# Exploring roles for essential proteins in yeast filamentous growth identifies the WASP homolog Las17 as a regulator of the Cdc42-dependent fMAPK pathway

**DOI:** 10.64898/2026.09.21.751561

**Authors:** Atindra N. Pujari, Ankita Priyadarshini, Zhijian Li, Ankita S. Darekar, Deanna Williams, Dale Climie, Sondra Bahr, Helena Friesen, Michelle Li, Joshua M. Oken, Robert M. Romito, Brenda Andrews, Charles Boone, Paul J. Cullen

**Author notes:** Corresponding author: Paul J. Cullen, PhD Address: 532 Cooke Hall, Department of Biological Sciences, State University of New York at Buffalo Buffalo, NY 14260-1300.

## Abstract

Cell differentiation into distinct cell types generates functional specialization in eukaryotic organisms. Many fungal species undergo filamentous growth, where cells differentiate into elongated and adhesive filaments capable of expansion and invasion into new environments. The budding yeast *Saccharomyces cerevisiae* also undergoes filamentous growth, and the regulatory pathways that control the response have been well characterized; however, the roles essential proteins play in this response have not been systematically explored. To address this gap in understanding, we constructed a collection of 332 temperature-sensitive (ts) alleles in 290 essential genes in a strain background that undergoes filamentous growth (called Σ1278b or Sigma). The ts Sigma collection showed differences in temperature sensitivity compared to a ts laboratory strain collection, revealing unexpected phenotypic diversity in essential alleles across populations of individuals. Screening the ts Sigma collection for phenotypes related to filamentous growth uncovered new phenotypes for >35% of essential alleles. New regulators of the Mitogen-Activated Protein Kinase (MAPK) pathway that regulates filamentous growth (fMAPK) were identified, including Las17, a homolog of Wiskott-Aldrich Syndrome Protein (WASP) in humans. Las17 regulated the fMAPK pathway by promoting delivery of the sensor protein, Sho1, and Rho GTPase Cdc42 to the plasma membrane. Las17 also functioned as a hub coordinating separate parallel aspects of the filamentation response. The widespread roles for essential proteins in regulating a eukaryotic differentiation response suggest broader roles for essential proteins than is currently appreciated.

**HIGHLIGHTS:**

- **A collection of conditional alleles of essential genes was generated in a yeast strain that undergoes filamentous growth**
  - 332 temperature-sensitive alleles in 290 essential genes were constructed and verified in the Sigma strain background
  - Conditional essentiality was compared between strain backgrounds, revealing unexpected phenotypic diversity in essential alleles between populations of individuals
- **The ts Sigma collection was screened for phenotypes in filamentous growth**
  - More than thirty-five percent of essential alleles showed a phenotype by at least one test
  - New cellular processes and protein functions were connected to the filamentous growth response
  - Together with analysis of nonessential genes, a more comprehensive picture of the filamentous growth response was produced genome-wide
- **New essential regulators of the fMAPK pathway were identified**
  - Sixteen new regulators in ten functional categories were identified
- **The WASP homolog Las17 regulates the fMAPK pathway**
  - Las17 controlled the localization and levels of the tetraspan protein Sho1 and Rho GTPase Cdc42
  - Functioned as a new regulator of plasma membrane delivery of cortical regulators
  - Acted as a hub that coordinated separate parallel regulatory inputs into the filamentous growth response

## INTRODUCTION

During cell differentiation, cells specialize into distinct cell types that perform specific functions. Cell differentiation creates a division of labor in multicellular organisms where specialized cells perform specific biochemical and morphogenetic functions. In humans, more than 200 distinct cell types have been identified, and new initiatives suggest the number of differentiated cell types to be much higher (Tabula Sapiens *et al*. 2022; Hatton *et al*. 2023; Yao *et al*. 2023; Rood *et al*. 2025). Regulatory programs control specialization into specific cell types, which can occur through intrinsic developmental programming or be induced by extrinsic cues. Evolutionarily conserved signaling pathways, including Hedgehog, Wnt, Notch, and Mitogen-Activated Protein Kinase (MAPK) pathways, and their cognate transcription factors, control cell-type specification by the induction of specific target genes (Peterson *et al*. 2022; Ding *et al*. 2026). These regulatory pathways function in interconnected networks that integrate multiple signals to produce a highly coordinated response. Inappropriate cross-talk within these networks can be disastrous for cells, leading to de-differentiation and diseases including cancers (Guo *et al*. 2020). A comprehensive understanding of the regulatory processes that control cell differentiation is therefore an emerging priority. The discovery of new cell types suggests the regulation of cell differentiation is considerably more complicated than is currently appreciated.

Most fungal species undergo cell differentiation to specific cell types. Depending on the species, specialized cells coordinate mating between haploid cells, sporulation of diploids, and nutrient foraging. Most fungal species can undergo filamentous growth, a fungal differentiation response composed of elongated and interconnected cells with unique adhesion properties (Gow and Lenardon 2023; Lorenz 2024). Filamentous growth facilitates expansion of mycelial cells outwards into new environments, and downwards into surfaces presumably to facilitate the acquisition of nutrients (Borneman and Pretorius 2015; Kiss *et al*. 2019). In many plant and animal fungal pathogens, filamentous growth is required for virulence (Lo *et al*. 1997; Borneman and Pretorius 2015; Woolford *et al*. 2016). Fungal species also form biofilms/mats that attach to surfaces and promote colonial expansion by surface spreading (Lagree and Mitchell 2017). Filamentous growth and biofilm/mat formation are related responses that utilize overlapping adhesion molecules and nutritional triggers. Accordingly, biofilm/mat formation is a hallmark of pathogenesis (Harding *et al*. 2009). The budding yeast *Saccharomyces cerevisiae* undergoes filamentous/invasive/pseudohyphal growth (Madhani and Fink 1998a), and biofilm/mat formation (Reynolds and Fink 2001), which provides a convenient model to understand how cells differentiate into specialized cell types (Kumar 2021). In *S. cerevisiae*, filamentous growth is typically studied in “wild” strain backgrounds, like the Σ1278b or Sigma background, because commonly used strains in research settings (e.g., S288c) have lost the ability to undergo filamentous growth due to genetic manipulation in the laboratory (Liu *et al*. 1996).

Genetic screens (Lorenz and Heitman 1998; Palecek *et al*. 2000), transposon screens (Mosch and Fink 1997; Jin *et al*. 2008), and genome-wide deletion collections constructed in strains that undergo filamentous growth (Ryan *et al*. 2012) have identified many of the regulatory proteins and pathways that control filamentous growth in *S. cerevisiae*. Like other eukaryotic differentiation responses, filamentous growth is regulated by regulatory pathways that operate in an integrated network (Borneman *et al*. 2006; Bandyopadhyay *et al*. 2010; Chavel *et al*. 2010; Johnson 2011; Bahar *et al*. 2023). Well-defined pathways that regulate filamentous growth include major nutrient control pathways like TOR (Cutler *et al*. 2001), Ras2-cAMP-protein kinase A (Ras-PKA) (Gimeno *et al*. 1992; Robertson and Fink 1998; Mosch *et al*. 1999), and the filamentation Mitogen-Activated Protein Kinase (fMAPK) pathway (Roberts and Fink 1994). These pathways are conserved across species and regulate morphogenetic responses from yeast to humans (Braicu *et al*. 2019). The fMAPK pathway is regulated by a mucin glycoprotein, Msb2 (Cullen *et al*. 2004), that operates at the plasma membrane with the tetraspan protein, Sho1 (O’rourke and Herskowitz 1998; Cullen *et al*. 2004), and the cysteine-rich protein, Opy2 (Wu *et al*. 2006). The mucin signaling complex regulates a canonical MAPK pathway that is controlled by the ubiquitous Rho GTPase Cdc42. Although the regulatory pathways that control filamentous growth have been highly studied, at many levels the regulation of filamentous growth remains mysterious, especially as new factors continue to be identified.

Most screens performed to identify new regulators of filamentous growth have focused on nonessential genes. By comparison, the diverse collection of essential genes, which cannot be deleted from cells, has not been systematically analyzed for roles in filamentous growth. Essential proteins perform key biological functions (Davierwala *et al*. 2005), and studies of conditional temperature-sensitive (ts) alleles and promoter-regulated genes/alleles (Mnaimneh *et al*. 2004; Breslow *et al*. 2008) have provided important insights into essential gene function (Hughes *et al*. 2000; Ben-aroya *et al*. 2008; Li *et al*. 2011). The collections of conditional essential alleles in yeast are not present in strains that undergo filamentous growth, which creates a gap in understanding the roles essential proteins play in this fungal differentiation response. To identify new roles for essential genes in the filamentous growth response, a collection of ts alleles from a laboratory strain (Li *et al*. 2011) was transferred to the Sigma strain background. When compared, the Sigma and S288c ts alleles showed a high degree of variation in conditional essentiality, indicating that ts phenotypes vary extensively between individuals. Evaluating the ts Sigma collection by tests for filamentous growth identified new roles for essential proteins and processes, with more than 35% of alleles showing phenotypes by at least one test. Secondary screens identified new regulators of the fMAPK pathway, including a role for the Wiskott-Aldrich Syndrome protein (WASP) homolog, Las17, in delivery of the sensor protein Sho1 and Rho GTPase Cdc42 to the plasma membrane. Given the high degree of functional conservation across eukaryotes (Wang *et al*. 2015), our study provides a roadmap for the roles essential proteins play in regulating cell differentiation across eukaryotes.

## RESULTS

### Constructing the Sigma ts collection and comparing conditional essentiality between individuals

A collection of ts alleles from a laboratory strain [S288c, (Li *et al*. 2011)] was transferred to a strain that undergoes filamentous growth [Sigma (Gimeno *et al*. 1992; Liu *et al*. 1996)]. Each ts allele and adjacent antibiotic-resistance cassette were amplified by polymerase chain reaction (PCR) from chromosomal DNA derived from the S288c ts collection. PCR products were introduced into the Sigma background by transformation, and antibiotic-resistant transformants were assessed for temperature sensitivity at 37°C and confirmed by PCR Southern analysis. In total, a collection of 332 ts alleles representing 290 unique genes was constructed (called the ts Sigma collection). The ts Sigma collection contains alleles for one-third of all essential genes (*Fig. S1*), representing most essential cellular functions (Winzeler *et al*. 1999; Giaever *et al*. 2002; Li *et al*. 2011).

Most genes essential in the S288c background are also essential in the Sigma background [**Fig. 1**, left 5% (Dowell *et al*. 2010)]. Compared to true essentiality, conditional essentiality driven by alleles has not been extensively explored. To explore conditional essentiality between individuals, the growth of Sigma ts strains was compared to the growth of the same alleles in S288c strains from (Li *et al*. 2011). Growth of ts strains was measured by assessing colony density by ImageJ analysis at 30°C, 32°C, and 34°C. Values were normalized to wild-type values and used to compare the growth rates of the same alleles at the same temperatures (*Table S3*). By this test, 36% of alleles exhibited similar growth profiles between strains, while 64% showed different growth patterns (**Fig. 1**, right). Most strains that showed different growth profiles grew better in the S288c background (80%), which may reflect selection for temperature sensitivity during construction of the collection. These findings can account for the limited recovery of alleles in the S288c background in the Sigma background (332 alleles of the 650 alleles attempted). These findings also highlight the broad variability in conditional essentiality between individuals, which show extensive genetic and phenotypic variation (Peter *et al*. 2018). Because conditional essentiality contributes to phenotypic variation across individuals, these findings may extend to understanding this aspect of allelic variation in other species.

**Figure 1.**
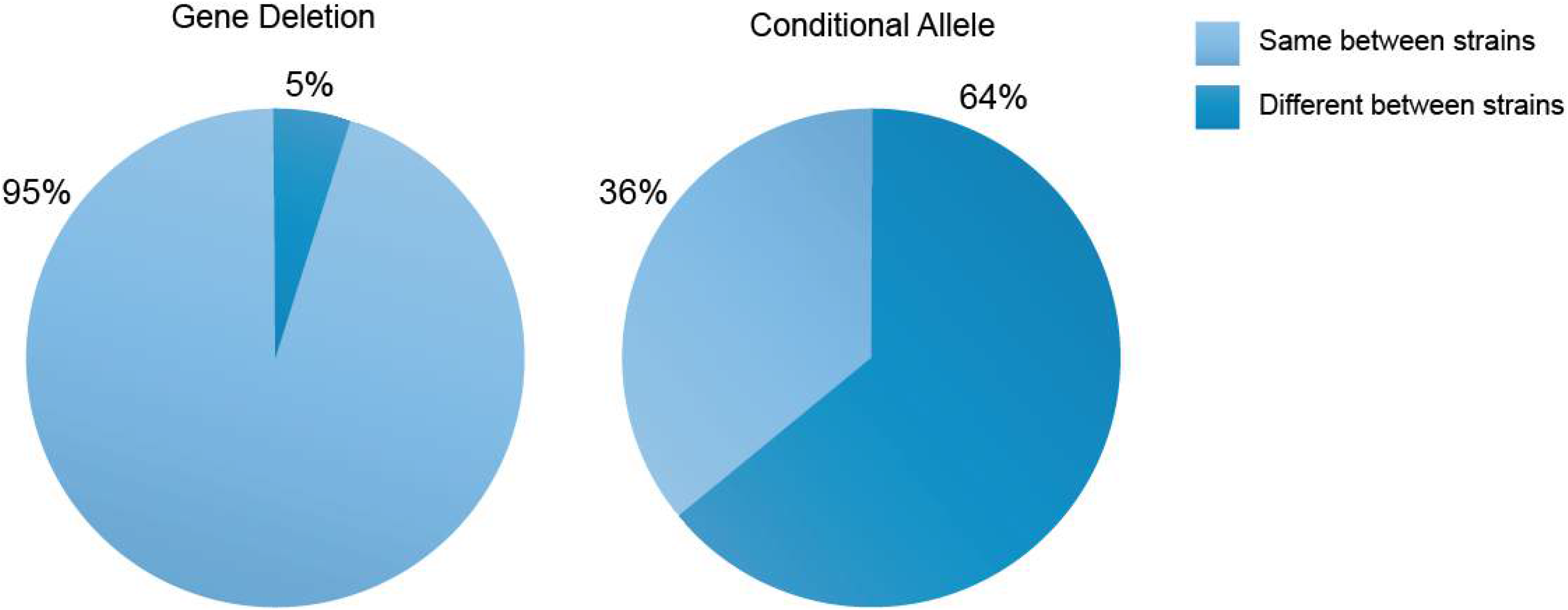
Differences in growth in essential genes and alleles between strain backgrounds. Pie charts showing similarities (light blue) and differences (dark blue) in growth between Essential Genes (left) and Conditional Alleles (right) in *S. cerevisiae* that were compared between the S288c and Sigma strain backgrounds at 30°C, 32°C, and 34°C. The raw data can be found in *File 1* (unwashed plates), and the analyzed growth intensity is shown in *Table S3*.

### Screening the Sigma ts collection for phenotypes in filamentous growth

We next screened the Sigma ts collection for phenotypes related to filamentous growth. Some ts alleles exhibited growth defects or tight ts phenotypes that interfered with the evaluation of filamentous growth. To address this caveat, only alleles that showed a growth rate of >50% compared to wild-type cells were considered for analysis. This cutoff underestimated the role of essential alleles in the filamentation program but reduced false positives. Each of the 332 strains was examined in three functional tests in separate biological replicates. The raw data from this comprehensive phenotypic survey is available (*File Set S1-S3*). *Table S4* summarizes the results of the phenotypic survey, and the collection is available as a community resource.

One aspect of filamentous growth is invasive growth. Invasive growth is measured by the plate-washing assay, where spots or patches of cells washed off semi-solid agar plates reveal an invasive scar (Roberts and Fink 1994). The plate-washing assay was performed in 96-well format by spotting cells onto YPD plates that were incubated at different temperatures (30°C, 32°C, and 34°C, *File Set S1*, see *Table S5* for key). Plates were incubated for 4d alongside controls that lacked (*ste12Δ*) or showed elevated activity (*dig1Δ*) of the fMAPK pathway (**Fig. 2A**, Invasion). Alleles with <50% growth rates compared to wild-type cells were excluded from the analysis (23 alleles [7% of all the alleles tested]; see *Table S3* for growth rate analysis). By the plate-washing assay, 28 alleles (8%) showed a strong defect in invasive growth (**Fig. 2A**, e.g., *myo2-14*), and 13 alleles (4%) showed hyper-invasive growth (**Fig. 2A**, e.g., *cdc37-1*). In total, 41 of the 332 alleles tested (12%) showed a phenotype in invasive growth by the plate-washing assay (*Table S4*).

**Figure 2.**
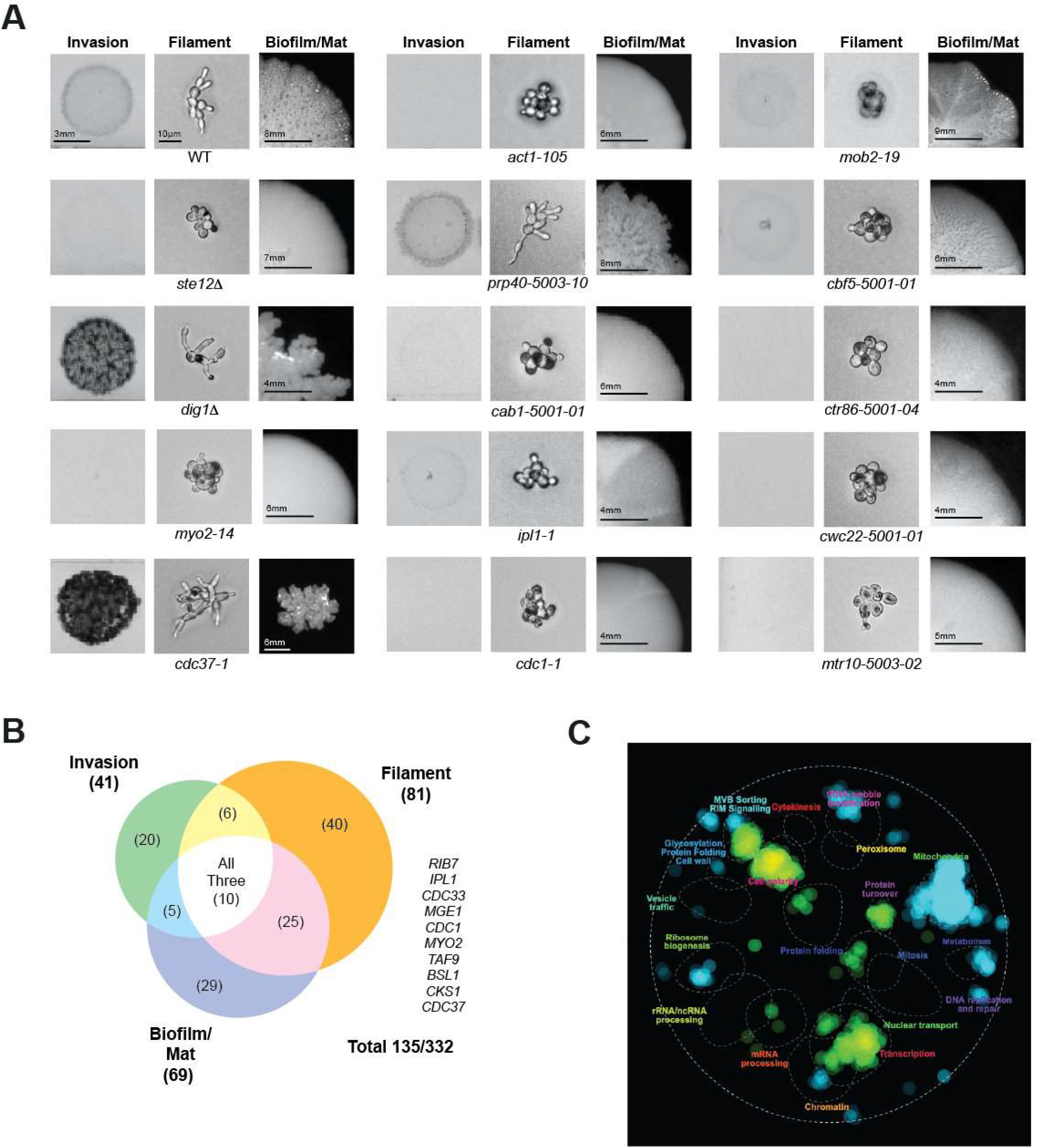
Phenotypes of ts essential alleles in filamentation assays. **A**) Examples of invasive growth (Invasion) by the plate-washing assay. Control cells and alleles were incubated at 32°C. Raw data can be found in *File Set S1.* Examples of filament formation (Filament) by the single-cell assay performed at 32°C. Raw data can be found in *File 2*. Examples of Biofilm/mat formation performed at 32°C. Examples can be found in *File 3.* Controls *ste12Δ* and *dig1Δ* are also shown. Scale bars are as indicated. **B**) Venn diagram. **C**) The Cell map. Blue, nonessential genes; Green, essential genes.

The ts alleles were next examined for filament formation by the single cell invasive growth assay (Cullen and Sprague 2000). Cells were grown in batches on glucose-limited semi-solid agar media at 30°C and 32°C (*File Set S2,* see *Table S6* for key), and filament formation was assessed by microscopic examination at 24 h. Eight alleles (2.4%) did not produce buds under this condition and were excluded from the analysis. These growth-defective alleles differed between tests, possibly due to the specific conditions tested. Microscopic examination showed that 74 alleles (22%) were defective for filament formation (**Fig. 2A**, Filament, e.g., *act1-105*), based on the failure to bud distally and produce elongated cells. By comparison, 7 alleles (2%) showed enhanced filament formation based on an increase in cell length compared to wild-type cells (**Fig. 2A**, e.g., *prp40-5003-10*). Overall, 81 of 332 alleles tested (24%) displayed a phenotype in filamentous growth by the single cell assay (*Table S4*).

Yeast also undergoes biofilm/mat formation, which can be assessed by a ruffled colony pattern as an indicator of surface adhesion (Reynolds and Fink 2001). Ts alleles were examined on low-agar (0.3%) media at 32°C to assess biofilm/mat formation (*File Set S3*, see *Table S7* for key). Pilot experiments identified the size cutoff at which ruffles form (*Fig. S2*), and four alleles (1%) fell below this cutoff and were excluded from the analysis. By this test, 54 alleles (16%) were defective for biofilm/mat formation (**Fig. 2A**, e.g., *cab1-5001-01*), and 15 alleles (4.5%) made biofilms/mats that were more wrinkly than wild type and were also defective for colonial expansion (**Fig. 2A**, e.g., *cdc37-1*). To account for the phenotypic diversity in biofilm/mat formation, a machine learning algorithm was employed. Models trained on controls showed phenotypic scoring that closely agreed with the original analysis, especially for the strongest hits (*Table S8*). Overall, 69 of 332 alleles tested (21%) showed a phenotype in biofilm/mat formation (*Table S4*). Summarizing these results, 191 phenotypic designations in 123 unique essential genes were uncovered by these tests. These findings demonstrate that filamentous growth and biofilm/mat formation are tightly connected to essential gene function.

We next compared the phenotypes of ts alleles in each of the filamentation tests to each other. Some alleles showed a phenotype by a single test (**Fig. 2B**, circles that did not intersect, *Table S9*). Other alleles showed phenotypes in two or three tests (intersecting circles). This pattern mirrored previous findings from the analysis of nonessential genes (Ryan *et al*. 2012). For alleles that showed phenotypes in multiple tests, most alleles had the same phenotypes in both (or all three) tests (*Table S10*). For example, 11/16 (68%) of the alleles strongly defective for invasion were also strongly defective for filament formation. A few alleles however showed opposing phenotypes. For example, *noc2-5001-01* was strongly defective for invasion but showed hyper-filamentous growth. Most alleles of a given gene also showed the same phenotype. For example, six different alleles of *PBR1* showed similar phenotypes across all three tests. In a few cases, allele-specific phenotypes were observed. The *las1-5004-01* and *las1-5001-05* alleles showed opposite phenotypes than the *las1-5005-01* and *las1-5006-01* alleles in filament formation. These results indicate that there is a strong functional overlap in genes with phenotypes in filamentous growth; however, test- and allele-specific phenotypes were also observed.

### Analyzing the roles essential proteins play in filamentous growth

We next examined essential genes that regulate filamentous growth by comparison to their known functions in the cell. Alleles defective in genes previously implicated in filamentous growth (12% or 15/124 genes tested, *Table S11*) were identified. These included components and regulators of the actin cytoskeleton [*act1-101, act1-105, pfy1-13, myo2-14, rho3-ser228, Table S11* (Cali *et al*. 1998)]. They also included regulators of signal transduction, like the Rho GTPase Cdc42 [*cdc42-1* (Mosch *et al*. 1996; Peter *et al*. 1996; Leberer *et al*. 1997)] and TOR pathways [*tap42-11* (Cutler *et al*. 2001)], regulators of the cell cycle [*cks1-13* and *cdc28-1* (Tobe et al. 2009)], chaperones that control protein folding [e.g., *cdc37-1* (Cowen and Lindquist 2005; Hawle *et al*. 2007; Yang *et al*. 2007; Shapiro *et al*. 2009; Jarosz and Lindquist 2010; Girstmair *et al*.2019; Hossain *et al*. 2021; Robbins and Cowen 2023)], and genes involved in protein translation [*DED1*, (Gilbert *et al*. 2007)]. Therefore, the tests uncovered known regulators of the filamentous growth program.

Roles for essential genes in filamentous growth were also identified that had not been previously reported (108/124 genes, 88% of the essential genes identified with filamentation phenotypes). A subset of alleles were defective in genes previously tied to pathogenic responses in other fungal species, like *rho1-td* (Corvest *et al*. 2013) and *pkc1-1* (Borah *et al*. 2011). However, most genes have not been previously tied to filamentous growth regulation in any species (61%, 76/124 new genes tested). These genes fell into several functional categories [*Table S12,* (Ashburner *et al*. 2000)]. Major functional categories included transcription, with subclasses in chromatin remodeling, mediator, transcription initiation factors/TFIID, and general transcription factors; mRNA regulation, including subclasses in mRNA capping, mRNA 3’ processing and polyadenylation, and pre-mRNA splicing; RNA regulation, including ribosome biogenesis, RNA degradation and surveillance, translation initiation, tRNA biosynthesis and modification; protein folding and turnover; and protein trafficking. A few genes comprised the functional categories of DNA replication, chromosome structure and cohesion, nuclear transport, protein lipid modification, metabolic biosynthesis, calcium signaling, and Fe-S cluster assembly. Four of the six alleles with phenotypes in splicing showed hyper filamentous growth, which indicates that splicing may function in an inhibitory capacity (*HSH49, SNU56, YHC1,* and *SPP381*). Therefore, the approach identified new genes that connect to the filamentous growth program in distinct functional categories.

The phenotypes of essential alleles were compared to the phenotypes of nonessential mutants from analysis of a non-essential deletion collection made in the Sigma background (Ryan *et al*. 2012). By mapping genes to specific processes (Usaj *et al*. 2017), essential and non-essential genes fell mainly into non-overlapping categories (**Fig. 2C**, *Table S13*). Compared to nonessential genes, where 1443 out of 3876 (37%) impacted filamentous growth (Ryan *et al*. 2012), 123 out of 290 essential genes (42%) showed phenotypes in filamentous growth. The high number of essential genes that showed phenotypes in filamentous growth may not be surprising because essential proteins have a high density of genetic interactions (Davierwala *et al*. 2005). In total, 1566/6000 (26%) of *S. cerevisiae* genes showed a phenotype in filamentous growth and/or biofilm/mat formation. This number may be an underestimate because not all genes were examined (4250/4889 nonessential and 290/1011 essential genes), and for the essential genes tested, only a single allele was examined.

### The Wiskott-Aldrich Syndrome Protein (WASP) homolog Las17 regulates the fMAPK pathway

Essential alleles may exhibit phenotypes in filamentous growth for many reasons, and it might not be surprising that cells with defects in actin function and morphogenesis, nutrient metabolism, transcription, and translation for example show phenotypes in filamentous growth. However, some essential proteins may play a regulatory role in filamentous growth. To identify essential genes that regulate the filamentous growth program, ts alleles with the strongest phenotypes were examined for altered activity of the fMAPK pathway, a central regulatory pathway of filamentous growth. The activity of a transcriptional reporter for the fMAPK pathway [*FRE-lacZ* (Madhani and Fink 1997)] was assessed in a panel of eighty three strains that showed phenotypes in filamentous growth and whose gene functions spanned distinct functional categories. Sixteen of the eighty-three alleles tested showed reduced *FRE-lacZ* activity corresponding to a reduction in filamentous growth by functional tests (**Fig. 3A**). The remaining essential alleles may regulate other signaling pathways or function through mechanisms not explored here.

**Figure 3.**
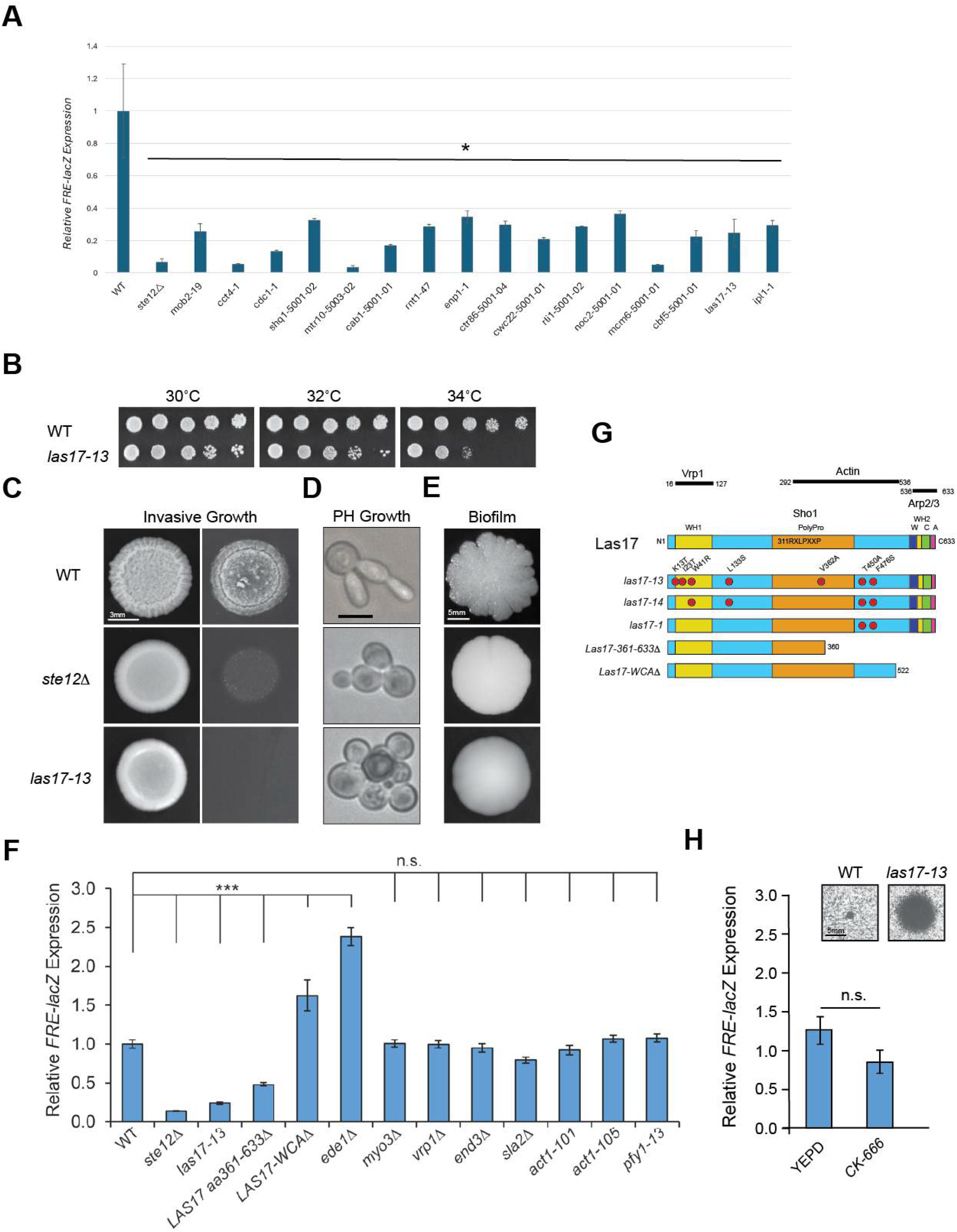
New fMAPK regulators include the WASP homolog, Las17. **A**) fMAPK pathway activity based on *FRE-lacZ* activity. Cells were precultured at 26°C in SD-URA media for 16 h. Then cells were incubated in YEPD at 30°C for 6 h. Experiments were performed in triplicate, and average values are shown with error bars representing the standard error of the mean. Asterisk, p-value <0.05. **B**) Serial dilutions of wild-type cells (WT, PC313) and the *las17-13* mutant were spotted onto YPD media at 30°C, 32°C, and 34°C. Plates were photographed at 4d. **C**) Invasive growth by the plate-washing assay of the *las17-13* mutant compared to wild-type cells and the *ste12*Δ control. Cells were grown for 3 d at 32°C. The plate was photographed, washed, and photographed again after washing. Bar, 3 mm. **D**) Examination of filament formation of the indicated strains in phenotypic assays at 32°C. Bar, 5 microns. **E**) Cells in panel 3B were grown on YPD media with 0.3% agar to observe formation of biofilms/mats. Bar, 5 mm. **F**) Evaluation of the indicated mutants for fMAPK pathway activity. The *FRE-lacZ* reporter on a plasmid was transformed into WT (PC313), *ste12*Δ (PC539), *las17-13* (TUBE #9), and *las17-14* (TUBE #10) mutants. β-galactosidase assays were performed at permissive and semi-permissive temperatures. Experiments were performed in three biological replicates. Error bars indicate standard error of the mean. **G**) Domain organization of the Las17 protein. Regions for interaction with Actin, Arp2/3 (WCA domain), Sho1, and Vrp1 have been established and are labeled and marked by black lines. Sites of mutations in *las17* alleles are represented by red circles and were determined by DNA sequencing analysis. Two C-terminal truncations are also shown. **H)** p*FRE-lacZ* activity of wild-type cells (PC313) grown in YEPD and exposed to CK-666. Experiments were performed in three biological replicates; error bars denote standard error of the mean. Inset, sensitivity of wild-type cells and the *las17-13* mutant to CK-666 after 2 d at 30°C. Bar, 5 mm.

One of the ts alleles that regulated the fMAPK pathway was defective for the Las17/Bee1 protein, which is a homolog of the human Wiskott-Aldrich Syndrome Protein [WASP **Fig. 3A**, *las17-13*]. The *las17-13* allele showed broad temperature sensitivity, which allowed a clear evaluation of its role in filamentous growth (**Fig. 3B**). The *las17-13* allele was defective for invasive growth (**Fig. 3C**), pseudohyphal growth (**Fig. 3D**), and biofilm/mat formation (**Fig. 3E**). Two alleles of *LAS17*, *las17-13* and *las17-14* showed similar defects in *FRE-lacZ* activity over a range of temperatures and fMAPK pathway-dependent reporters (**Fig. 3F**, *Fig. S3C*). Collectively, these results indicate that Las17 functions as a positive regulator of the fMAPK pathway, representing the first connection between WASP and MAPK pathway regulation in any system.

We first tested whether Las17 regulates the fMAPK pathway through its established function in the cell. WASP proteins from yeast to humans nucleate branched actin filaments (Li 1997; Madania *et al*. 1999; Winter *et al*. 1999; Derry *et al*. 2018) that function in endocytosis and other aspects of protein trafficking (Galletta and Cooper 2009; Ghose *et al*. 2022). To define whether Las17 regulates the fMAPK pathway by this function, alleles defective for actin function were examined. Alleles in the actin gene (*ACT1*) did not show reduced *FRE-lacZ* activity (**Fig. 3F** and *Fig. S4A*). These alleles were defective for filamentous growth (*Fig. S4B*), as has been reported (Cali *et al*. 1998). Las17 is a multidomain protein that nucleates actin polymerization through the Arp2/Arp3 complex (**Fig. 3G**). Conditional alleles of *arp2/arp3* were inviable at all temperatures tested, so the pharmacological inhibitor CK-666 was examined (Nolen *et al*. 2009). CK-666 showed hypersensitivity in the *las17-13* mutant, which supports the idea that this inhibitor specifically targets the Arp2/3 complex (*Fig. S4C*). CK-666 did not impact *FRE-lacZ* activity (**Fig. 3H** *Fig. S4D*). Likewise, a version of Las17 lacking the C-terminal WASP homology II, central, acidic (WCA) domain, which interacts with Arp2/3 (Rohatgi *et al*. 1999; Goode *et al*. 2015) showed normal fMAPK pathway activity (**Fig. 3, F-G,** *LAS17-WCA*Δ). The N-terminal region of Las17 binds to Verprolin, which also regulates Arp2/3 activity (Lechler *et al*. 2001; Sun *et al*. 2017); however, loss of verprolin did not impact fMAPK pathway activity (**Fig. 3, F-G**, *vrp1*Δ). Therefore, Las17 regulates the fMAPK pathway outside of the Arp2/3 complex. Las17 also directly interacts with actin [**Fig. 3G**, (Higgs *et al*. 1999; Rohatgi *et al*. 1999; Marchand *et al*. 2001; Goley and Welch 2006; Alekhina *et al*. 2017; Narvaez-ortiz *et al*. 2024)]. Deletion analysis identified a region in the C-terminus (Las17-361-633Δ) that lacks a proline-rich domain that binds actin (Urbanek *et al*. 2013), as being required for fMAPK pathway activity (**Fig. 3F**).

A central function for Las17-dependent actin nucleation is the regulation of endocytosis (Johansen *et al*. 2016; Luan *et al*. 2018; Hummel and Kaksonen 2023). Cells defective for endocytosis were not defective for fMAPK pathway activity (**Fig. 3F**, *end3*Δ, *vrp1*Δ, and *sla2*Δ; *ede1*Δ showed elevated fMAPK activity). Most of these mutants showed defects in invasive growth (*Fig. S5*), which indicates that endocytosis has a functional connection to filamentous growth through another mechanism. Altogether, these results indicate that Las17 regulates the fMAPK pathway by a mechanism that is separate from its essential function in the cell. A moonlighting function for Las17 is consistent with evidence that the *las17-13* mutant showed defects in filamentous growth at temperatures that did not compromise cell viability.

### Las17 regulates the level and localization of the tetraspan protein Sho1 and Rho GTPase Cdc42

To determine how Las17 regulates the fMAPK pathway, we defined at which point in the fMAPK pathway the protein functions. The fMAPK pathway requires a subset of proteins that also regulate the mating pathway [**Fig. 4A**, Cdc24, Bem1, Cdc42, Ste20, Ste11, Ste50, Ste7, and Ste12 (Madhani and Fink 1998b; Bardwell 2006; Saito 2010)]. Las17 was not required for growth arrest in response to the mating pheromone α-factor (**Fig. 4B**), which indicates that Las17 acts on a protein that does not function in the mating pathway (Msb2, Sho1, Opy2, Bem4, Kss1, and Tec1). We next performed genetic epistasis analysis, by examining suppression of the signaling defect of the *las17-13* mutant with gain-of-function alleles that hyperactivate the fMAPK pathway. These results showed that Las17 functions between Msb2 and Ste11 in the fMAPK pathway (*Fig. S6*), which include three proteins, Opy2, Sho1, and Bem4.

**Figure 4.**
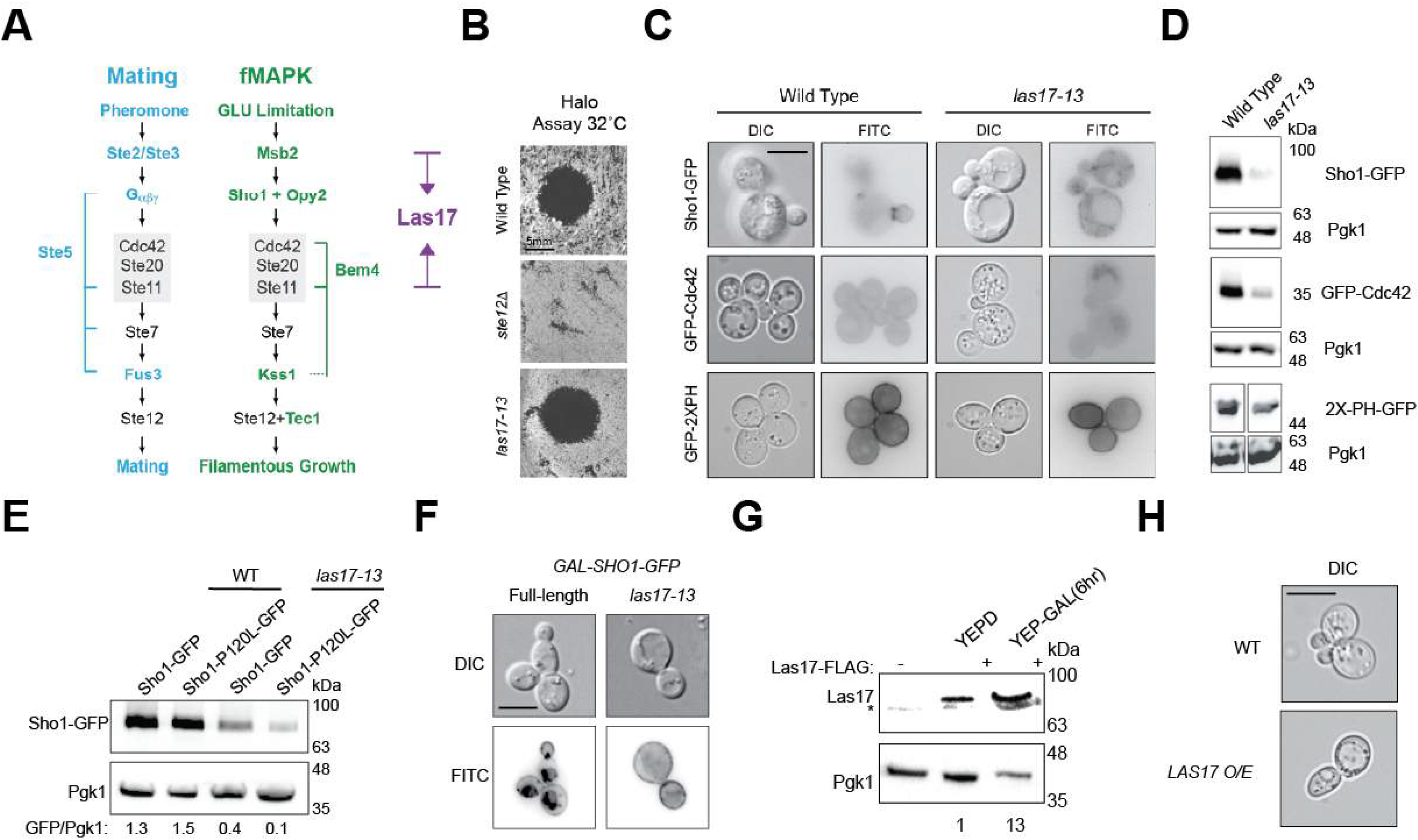
Las17 impacts the levels and localization of fMAPK pathway sensor Sho1 and GTPase Cdc42. **A**) The fMAPK and mating pathways share components (black) and contain pathway-specific factors (blue, fMAPK; green, mating). Purple designates putative sites of Las17 function based on the genetic suppression analysis presented in *Fig. S6.* **B**) Halo assay. Equal concentrations of WT (PC313), *ste12*Δ (PC539), and *las17-13* cells were spread onto YPD media. The mating pheromone α-factor at a concentration of 1 mg/ml was spotted in 5 μL (left) and 10 μL (right) aliquots to observe formation of halos. Pictures were taken after 2 d of growth at 30°C and 32°C. Bar, 5 mm. **C**) The localization patterns of pSho1-GFP, pGFP-Cdc42, and pGFP-2XPH-PLCδ were examined in YPD media in wild-type cells and the *las17-13* mutant. Bar, 5 microns. **D**) Levels of pSho1-GFP, pGFP-Cdc42, and pGFP-2XPH-PLCδ assessed by IB analysis. GFP-tagged proteins were detected with monoclonal mouse anti-GFP antibodies. Antibodies to Pgk1 were used as a control for total protein levels. **E)** Immunoblot analysis measuring hyperactive Sho1 levels. Extracts from wild-type [WT, (PC313)] and the *las-17-13* mutant harboring pRS316 as a control for pSHO1^P120L^-GFP. **F)** Localization of Sho1-GFP in a pulse-chase experiment after a 2h induction in YEP-GAL (2%) followed by chase in YEPD medium for 3h at 32°C. Bar,10 microns. **G)** Immunoblot analysis measuring FLAG-tagged Las17. Pgk1, loading control. **H)** Representative cells showing wild-type cells (WT) or cells carrying a high-copy Las17 (*LAS17 OE).* Grown to mid-log phase. Bar, 5 microns.

We next examined the localization and protein levels of fMAPK pathway specific proteins in the *las17-13* mutant. Functional green-fluorescent protein fusions were examined by fluorescence microscopy and immunoblot analysis using anti-GFP antibodies. We observed that the localization of Sho1-GFP, which is normally enriched in buds and the mother-bud neck (Raitt *et al*. 2000; Reiser *et al*. 2000), was mis-localized in the *las17-13* mutant, being found in a diffuse pattern (**Fig. 4C**). The levels of the Sho1-GFP protein were also reduced in the *las17-13* mutant compared to wild-type cells (**Fig. 4D**, >3-fold). The conserved Rho GTPase Cdc42 is a master regulator of cell polarity and signaling. Although Cdc42 regulates the fMAPK and mating pathways (Bardwell 2006; Saito 2010; Ma and Nicolet 2023), the protein is maintained at high levels to permit fMAPK pathway activity (Gonzalez and Cullen 2022). A functional GFP-Cdc42 fusion protein was also mis-localized in the *las17-13* mutant (**Fig. 4C**, GFP-Cdc42) and like Sho1-GFP, GFP-Cdc42 was found at lower levels in the cell (**Fig. 4D**, >2.8-fold). These proteins stood out compared to Msb2-GFP, found enriched in vacuoles, and Opy2-GFP, found at the plasma membrane (*Fig. S7B*). As a control, the localization (*Fig. S7D*) and levels (*Fig. S7E*) of GFP-Ste20, an effector of Cdc42 and shared component between the mating and fMAPK pathways (Peter *et al*. 1996; Leberer *et al*. 1997; Moran *et al*. 2019), showed normal levels and localization in the *las17-13* mutant. Las17 might impact the targeting of these proteins by influencing the levels of phosphatidylinositol (4,5)-bisphosphate [PI(4,5)P2], a mark for targeting proteins to the plasma membrane. To test this possibility, a pGFP-2XPH-PLCδ reporter was evaluated by localization and levels in the cell (Stefan *et al*. 2002). By these tests, Las17 did not impact the levels of PI(4,5)P2 at the plasma membrane (**Fig. 4D**). These results demonstrate that Las17 regulates the levels and localization of proteins that govern the activity of key proteins that regulate the fMAPK pathway.

### Las17 regulates the delivery of Sho1 and Cdc42 to the plasma membrane under conditions that favor filamentous growth

In addition to endocytosis of vesicles from the plasma membrane, Las17 also regulates vesicular trafficking between compartments (Chang *et al*. 2003; Takenawa and Suetsugu 2007). Thus, the localization of Sho1 was examined in protein trafficking mutants. Sho1-GFP accumulated at the plasma membrane in endocytosis mutants (*Fig. S5A*, for *end3*Δ). Because Las17 is expected to be required for endocytosis of Sho1 from the plasma membrane, we conclude that Sho1 fails to be delivered to the plasma membrane in the *las17-13* mutant. Failure of Sho1 and Cdc42 to be delivered to the plasma membrane may result in its degradation by the secretory pathway, which would account for the reduced Sho1 levels in the *las17-13* mutant.

To better understand Sho1 trafficking in the cell, protein levels were controlled by an inducible promoter (p*GAL1-Sho1-GFP*) that when shut off by growth in glucose allowed temporal assessment of Sho1 location by pulse chase. Sho1-GFP localized at the plasma membrane at early time points (30 min) and then accumulated in the pre-vacuolar compartment (3h-5h) in wild-type cells (**Fig. 4F**). By comparison, Sho1-GFP was localized at the plasma membrane in the *las17-13* mutant throughout the course of the experiment (**Fig. 4F**). Therefore, overexpression of Sho1 was able to partially bypass the plasma membrane localization defect of the *las17-13* mutant, but once there, the protein was retained at that site. Collectively, these results show that Las17 contributes a new aspect of Sho1 regulation, by regulating delivery of the protein to the plasma membrane.

We next asked whether Sho1 is regulated by Las17 in its activated state. Gain-of-function variants of Sho1 have been identified that show hyperactivity of the fMAPK and HOG pathways (Tatebayashi *et al*. 2007; Vadaie *et al*. 2008). Two of these variants, Sho1^P120L^ -GFP and Sho1^S220F^-GFP, also showed reduced levels in the *las17-13* mutant, indicating that Las17 is required to maintain active levels of the fMAPK pathway (**Fig. 4E**, *Fig. S6C*). It was previously shown that the poly-proline domain of Las17 interacts with Sho1 (Tong *et al*. 2002b), but a region of Las17 containing the poly-proline domain (310-RXLPXXP, best match) was not required for Sho1 localization or fMAPK pathway activity (**Fig. 3F-G**). Las17 was not required for the expression of the *SHO1* gene (*Fig. S7A*), despite being required for Sho1-GFP localization (*Fig. S7B*). Thus, Las17 may directly bind to Sho1, or indirectly by regulating the trafficking of Sho1, to regulate its levels or localization in this context. Finally, we tested whether Las17 may be regulated during filamentous growth. The level of Las17-FLAG was higher under filamentous growth conditions (**Fig. 4G**). Similarly, high levels of Las17, expressed on a high-copy plasmid, stimulated filamentous growth (**Fig. 4H**). These results provide a mechanism by which Las17 can selectively augment the activity of the filamentous growth pathway in a condition-specific manner that is separable from its essential function.

### Las17 has separate parallel functions in regulating filamentous growth

The invasive growth defect of the *las17-13* mutant was more severe than the defect seen in cells completely lacking an intact fMAPK pathway (**Fig. 3C**, compare *las17-13* to *ste12*Δ). Thus, Las17 may have roles in regulating filamentous growth beyond MAPK pathway regulation. Las17 regulates endocytosis (Larriba *et al*. 1993; Naqvi *et al*. 1998; Feliciano and Di Pietro 2012), which, as shown above, was required for invasive growth independent of the fMAPK pathway (*Fig. S5A*). Endocytosis also contributes to the polarized growth of cells (Wu and Jiang 2005; Duan *et al*. 2021). We therefore hypothesized that Arp2/3 regulation of endocytosis may impact the polarized growth of filamentous cells. Consistent with this possibility, the Arp2/3 inhibitor, CK-666, inhibited filament formation (*Fig. S8A*) independent of the fMAPK pathway (**Fig. 3H**, *Fig. S4D*). Similarly, overexpression of *SHO1* causes hyper-polarized growth outside the fMAPK pathway (Pitoniak *et al*. 2015), which was also dependent on Las17 (*Fig. S8*, B and C*, las17-13*), the actin cytoskeleton (*Fig. S8D, act1-101, act1-105,* and *pfy1-13*), and the SH3 domain of Sho1 (*Fig. S8*). Endocytosis is also required for bud-site selection (Tuo *et al*. 2013), which in haploid cells involves a switch from axial to distal-unipolar budding to promote filament formation. The *las17-13* mutant exhibited bud-site-selection defects (*Fig. S8E*), which would be expected to interfere with filament formation. Bud-site-selection defects also cause a defect in fMAPK pathway activity (Basu *et al*. 2016); however, the levels of Sho1 and other proteins were not reduced in bud-site-selection mutants, indicating that Las17 and polar landmarks regulate the fMAPK pathway by separate mechanisms. Las17 was also required for cell polarization during mating (*Fig. S8D*). Therefore, Las17 functions as a hub coordinating separate parallel aspects of the filamentous growth response (*Fig. S8F*).

## DISCUSSION

Essential proteins are required for some of the most important and diverse cellular functions, yet due to their critical roles, they have not been systematically examined for their roles in cell differentiation and signal transduction. Here, we report the construction of a collection of conditional ts alleles of essential genes in a yeast strain background that undergoes a classical fungal differentiation response called filamentous growth. Although we succeeded in transferring nearly half the ts alleles from one strain background to another, many alleles did not recapitulate ts phenotypes, revealing differences in conditional essentiality between strains. Like most species, yeast strains from different niches contain an assortment of alleles that result in phenotypic differences (Peter *et al*. 2018) including drug sensitivity (Perlstein *et al*. 2007), which can be adaptive for survival (Vazquez-garcia *et al*. 2017). Different yeast strains also show differences in essentiality (Dowell *et al*. 2010), which can arise due to single or multiple differences between individuals (Hou *et al*. 2019). Likewise, human individuals possess extensive genetic variation that can lead to different phenotypes due to changes in gene expression (Yan *et al*. 2002) and protein function (Jordan *et al*. 2010). Much is known about essential genes in the human genome (Wang *et al*. 2015). For example, the loss of heterozygosity in essential genes underlies cancer vulnerabilities (Nichols *et al*. 2020). However, less is known about allelic variation in essential genes across individuals, which has the potential to contribute to many aspects of human health, drug sensitivity, and phenotypes. Our results in yeast indicate that allelic variation in essential genes likely represents an important source of phenotypic variation between individuals.

The Sigma ts collection was also screened for phenotypes related to filamentous growth. This revealed a prominent role for essential proteins in the growth response. Nearly forty percent of essential alleles tested showed a phenotype by at least one test. As a result of these screens, new roles for essential processes can be attributed to the filamentous growth response. Combined with a study to identify roles for nonessential proteins in filamentous growth (Ryan *et al*. 2012), this study provides a first approximation of the genome-wide contribution of a eukaryotic differentiation response, with approximately 25% of the genes being involved. Most essential genes in yeast are highly conserved throughout eukaryotes. Therefore, new roles for essential proteins in cell differentiation may apply to differentiation responses in other species. Fungal-specific essential genes that regulate filamentous growth may be suitable targets for combatting fungal pathogenesis.

Essential proteins were identified that regulate one of the main pathways that controls filamentous growth. One of these was the WASP homolog Las17 (Symons *et al*. 1996; Li 1997)] as a pathway-specific regulator of the fMAPK pathway (see *Fig. S9* for a model). This discovery represents the first connection between WASP and MAPK pathway regulation in any system. In humans, WASP nucleates branched actin formation and is a major regulator of endocytosis (Derry *et al*. 1994; Karpova *et al*. 2000; Tong *et al*. 2002a). In line with the general role of Las17 in controlling aspects of protein trafficking, Las17 was required for full levels and proper localization of the tetraspan protein Sho1 and Rho GTPase, Cdc42. Las17 interacts directly with the SH3 domain of Sho1 (Tong *et al*. 2002b) and may block sites of ubiquitination, as described for Cdc42 (Gonzalez and Cullen 2022). Alternatively, as an established regulator of endocytosis, Las17 may control the delivery of Sho1 and/or its recycling to the plasma membrane as has been described for Msb2 (Pitoniak *et al*. 2015; Gonzalez and Cullen 2022). In *C. albicans*, the Las17 homolog Wal1 plays a similar role in protein trafficking based on mutant phenotypes (Borth et al. 2010) and phenotypic profiling with chemical inhibitors (Douglas *et al*. 2009; Bar-yosef *et al*. 2017; Lash *et al*. 2023). The *C. albicans* WASP homolog Wal1 is required for the polarization of hyphal cells through interactions with the actin cytoskeleton (Walther and Wendland 2004; Borth *et al*. 2010; O’meara *et al*. 2016). Higher levels of sensor proteins resulting from Las17 function would be expected to sustain fMAPK pathway activity.

WASP/Las17 also controls protein trafficking from internal compartments, including the endosome (Chang *et al*. 2003). Moreover, WASP-related proteins, like Wiskott Aldrich Syndrome protein and SCAR homolog (WASH) participates in recycling proteins back to the plasma membrane through retromer (Gomez and Billadeau 2009; Guo *et al*. 2024). Yeast does not have a WASH homolog, and Las17 may regulate aspects of protein trafficking performed by WASH in higher organisms. Las17 also regulates at least two other aspects of filamentous growth, cell polarization and bud-site-selection, by parallel mechanisms. Finally, protein trafficking regulation of signaling can be highly variable across species. In the filamentous rice pathogen, *Magnaporthe oryzae*, the endosomal system acts as a docking platform for components of the homologous MAPK cascade to control the filamentation response (Wang *et al*. 2025).

In summary, essential proteins probably play a more prominent role in eukaryotic cell differentiation than is currently appreciated. Given the extensive conservation of essential protein functions across eukaryotes, essential proteins may have roles in regulating cell-type specification through conserved mechanisms.

## MATERIALS AND METHODS

### Yeast Strains, Media, and Growth Conditions

Unless otherwise indicated, haploid strains of *S. cerevisiae* come from the Σ1278b background (Liu *et al*. 1993). Yeast strains are listed in *Table S1*, and plasmids are listed in *Table S2*. Cells were grown at 30°C unless otherwise indicated. YPD, yeast extract (1%), peptone (2%), and dextrose (2%), or synthetic complete dextrose (SD) media containing 0.67% yeast nitrogen base (Sigma-Aldrich, St. Louis, Missouri) and 2% glucose were used for growth of yeast cells. S-Gal media was made by substituting 2% Galactose for glucose in SD media. Amino acids were added to the SD and S-Gal media as required. +AA denotes media containing all amino acids, -HIS denotes media containing all amino acids without histidine, and -URA denotes media containing all amino acids and lacking uracil. Strains were grown at a range of temperatures including 22°C (ambient temperature), 26°C, 30°C, 32°C, 34°C, 35°C, 36°C, and 37°C. To evaluate *HIS-*based growth reporters, the inhibitor 3-Amino-1,2,4-triazole [3-ATA (Struhl and Davis 1977)] was added to final concentrations of 2.5 mM, 5 mM, and 10 mM as indicated. CK-666 was used at a final concentration of 33.3 mM. Yeast strains were manipulated by standard techniques. Gene deletions, epitope tagging, and *GAL1p* promoter inductions strains were made by polymerase chain reaction (PCR) based strategies as described (Longtine *et al*. 1998). Some plasmids were used that came from a global overexpression collection (Gelperin *et al*. 2005).

To make strains using antibiotic-resistance marker, the antibiotic resistance cassettes, *KanMX6* and *HphMX4*, were PCR amplified and integrated at the gene locus as described (Goldstein and Mccusker 1999). The antibiotic geneticin sulfate (G418, Adipogen, CAS 108321-42-2) was added to YPD media to a final concentration of 0.36 mg/mL. Hygromycin was added to YPD media at a final concentration of 30 mg/ml.

To construct the *MAT***a** *ura3-52 leu2*Δ *his3*Δ (PC7693) strain, the parent Σ1278b strain PC313 (*MAT***a** *ura3-52*) was transformed with the pop-in pop-out HA-*URA3-*HA cassette (Schneider *et al*. 1995) amplified by primers to direct integration at the *LEU2* locus by homologous recombination. Positive *URA* ^+^*leu* ^-^ transformants were grown on media containing 5-floroorotic acid (5-FOA) to force out the cassette leaving an unmarked gene deletion. Similarly, the *ura3-52 leu2*Δ strain (PC7693) was transformed with the *HA-URA3-HA* cassette using primers targeting the *HIS3* locus, followed by growth on 5-FOA media to yield the *ura3-52 leu2*Δ *his3*Δ strain (PC7711). Plasmid p416GPD-LAS17 *URA3* was provided by the Breitenbach laboratory (Weber *et al*. 2021). *pCDC12-GFP::KanMX6* (PC7143) was constructed from YCplac181 Cdc12-GFP (Fares *et al*. 1996), by integration of the KanMX6 marker. DNA sequencing analysis was performed by Plasmidsaurus (https://plasmidsaurus.com/).

### Transferring TS alleles to the filamentous Σ1278b strain background

The ts alleles linked to the antibiotic-resistance marker (*KanMX6*) generated in the S288c background (Li *et al*. 2011) were moved to the Σ1278b background, which undergoes filamentous growth (Liu *et al*. 1993). The ts alleles and associated antibiotic resistance cassettes (*KanMX6*) were amplified by polymerase chain reaction (PCR) using the same forward (F1) and reverse (R4) primers used to construct the original ts collection. For a subset of ts alleles, PCR amplification was performed using primers pairs located ∼200 nucleotides (nts) upstream of the ATG start codon of the ORF and ∼200 nts downstream of the stop codon of the *KanMX6* gene. A collection of ts alleles with multiple isolates was generated in ten 96-well plates. Upon removal of duplicates, the final set of alleles was arrayed into 96-well plates. PCR amplified ts alleles and cassettes were transformed into a wild type Σ1278b strain (PC313). Transformants were spread on YPD+G418 semi-solid agar media to select for G418-resistant colonies. Transformants were transferred to YPD+G418 plates by replica plating and incubated at 26°C, 34°C, and 36°C to identify temperature sensitive colonies. Colonies defective for growth at elevated temperatures were streaked onto YPD+G418 along with wild-type cells at 26°C, 34°C, and 36°C as a control.

The ts alleles were verified by PCR Southern analysis. Genomic DNA was prepared as described (Pujari and Cullen 2024). Forward primers were designed ∼ 200 nts upstream of the F1 primer (*Table S2*). The same reverse primer was used that hybridized to the *TEF1* promoter within the *KanMX6* cassette. Isolates whose chromosomal DNA showed the expected product were frozen as -80°C stocks. Plasmid complementation was also used to confirm a subset of the ts strains. Two plasmid libraries were used that contained wild-type ORF on a CEN-ARS-based plasmid MoBY ORF collection or a 2.0 version in 2 micron plasmids (Ho *et al*. 2009). Strains were arrayed in 96-well format along with controls, and ∼100 strains were tested for complementation with the plasmid libraries. All strains tested passed plasmid complementation, confirming allele identity. For most alleles, multiple isolates were carried forward in the primary screens. In cases where the phenotype of one isolate did not match the phenotype of another, the slow-growing isolate was considered unless otherwise indicated to minimize phenotypes resulting from suppressors (Hawthorne and Mortimer 1963; Rasse-messenguy and Fink 1973; Liebman and Sherman 1976), which can benefit fitness by rescuing growth defects (Hou and Schacherer 2017).

### Construction of the Sigma ts collection

Strains were manipulated in batch by pinning with a 96-well pinning tool (VP Scientific, Catalog #VP408) sterilized by incubation in a bleach bath (50%) followed by autoclaved distilled deionized water, 70% ethanol, and two 95% ethanol baths. The pinning tool was flamed for sterilization and allowed to cool for 30 sec prior to use. A collection of 648 strains containing multiple isolates of 332 alleles was frozen as individual strains in separate tubes, designated by tube number, following confirmation by temperature sensitivity and PCR Southern analysis. To remove multiple isolates and include additional controls, a collection of 332 strains and controls were individually picked onto four 96-well plates, representing the final collection.

### CRISPR/CAS9 strategy to move select alleles

Some alleles were generated using a CRISPR/CAS9 approach (Jinek *et al*. 2012). sgRNA encoding target sequences for the CAS9 enzyme were subcloned into a pCAS9 plasmid [Addgene, (Ryan and Cate 2014)]. The homology-directed repair template was amplified by PCR using one forward and two reverse primers as described, containing the PAM site change and the desired mutations. To generate the *mrs6-2* allele (Bialek-wyrzykowska et al. 2000) a PCR product was generated to introduce changes resulting in two amino acid substitutions at positions S335P and G227V. Both mutations were checked by PCR amplification and DNA sequencing. DNA sequencing analysis was performed. The *las17-13* alleles also showed several substitutions (K13E, I23T, W41, L133S) in the N terminal of the protein.

### Comparison of growth rates of conditional alleles between strain backgrounds

Arrayed strains were unfrozen in 96-well format by pinning and incubated for 3d at 26°C. Alleles were transferred in 96-well cell culture plates (Corning Incorporated [#3596]), containing 200 uL of YPD + G418 in liquid media, and incubated for 15h at 26°C. Optical Density (OD) at A_600_ was measured by a 96-well spectrophotometer. Cell concentrations were adjusted to an OD of 1.0 by dilution in water. Cells were spotted onto YPD + G418 OmniTrays in 96-well format in 3 µL aliquots by a multichannel pipettor. Cells were periodically resuspended by pipetting to measure the OD and prevent settling. Two wild-type strains (PC313 and PC538), an invasion-defective strain [*ste12*Δ (PC539)], and a hyper-invasive strain [*dig1*Δ (PC3039)] were included as controls on each plate. Controls were transformed with the *pCDC12-GFP::KanMX6* (PC7143) plasmid to allow growth on YPD+G418 media.

Plates were spotted in replicates and incubated for 4d at 30°C, 32°C, and 34°C. Images were obtained with the Bio-Rad Molecular Imager ChemiDoc XRS+ (Catalog #170-8265) and Image Lab Software (https://www.bio-rad.com/en-us/product/image-lab-software) with zoom setting of 13.0 x 9.7. To determine growth rates, images were quantitated by ImageJ (FIJI) software (https://imagej.net/software/fiji/). Images were inverted, and the background signal was removed with the *Subtract Background* function with a rolling ball radius set to 20 pixels and the light background option enabled. Images were then analyzed by the Protein Array Analyzer tool (http://image.bio.methods.free.fr/dotblot.html). In the analysis settings, background subtraction was set to *None*, with the default rolling ball radius. A measurement grid corresponding to the 96-spot plate format was generated. The grid was aligned to colonies by selecting reference points to mark regions of interest (ROIs) across the plate. ROI size was adjusted for each image. Integrated density values for each position were measured and exported for analysis. The colony intensity for at least three wild-type controls was averaged, and outliers were removed. To obtain the relative growth rate, the average colony intensity of wild-type colonies was subtracted from the average colony intensity measured for each allele. Growth comparisons were made to the S288c background, which was performed in the same way except cells were grown in liquid culture (Li *et al*. 2011). Pilot experiments showed similar growth profiles between liquid and semi-solid agar media for a subset of alleles tested.

### Plate-washing assay

The plate-washing assay was performed as described (Roberts and Fink 1994). Pilot experiments were performed at a series of temperatures and time points to determine the optimal conditions for invasive growth. Invasive growth was examined at 4d at 30°C, 32°C, and 34°C. Each plate was assessed in biological replicates. Arrayed strains were spotted as described above. For washing, each part of the plate was rinsed in a stream of water at a flow rate of 7 ml/sec. Colonies were rubbed off plates by hand, with care taken not to damage the agar. To eliminate growth defects that could interfere with the assessment of invasive growth, an additional parameter was applied to the growth rate measurements determined above. Specifically, a threshold value was determined for colonies showing a growth defect at 30°C (55,000). An additional scoring criterion was applied above the base threshold, set to 50% of wild-type growth rate (65,000). Colonies with values below the threshold were excluded from analysis as growth defects could interfere with assessment of the invasive growth response. Colonies with values below the threshold at 32°C and 34°C were examined at 30°C for invasive growth. Some alleles with growth defects were incubated for extended time points (10d) before washing alongside controls.

### Single Cell Invasive Growth Assay

The single cell assay was performed as described (Cullen and Sprague 2000) with the following modifications. The ts mutants along with wild type strains (PC313) and the *ste11*Δ mutant (PC611) were inoculated into 5 ml of SD+AA media and grown for 16h at 30°C. The following day, 1 mL of cells were harvested by centrifugation, washed twice in sterilized water, and resuspended in 1 mL of sterilized distilled water. Approximately 50 μl of resuspended cultures were diluted in 1 mL of sterilized distilled water, and 5 μl of diluted cultures were spotted onto S+AA (lacking glucose) semisolid agar media. About 20 strains were spotted per plate equidistant from each other, and the plates were incubated at 32°C for 8-10 h (4-5 cell division cycles) to observe filamentous growth. Plates were removed from the incubator, equilibrated at 25°C temperature, and cells were observed my microscopy at 20X magnification. Alleles that failed to undergo budding under the conditions tested were excluded from analysis.

### Biofilm/Mat Assays

Biofilm/mat assays were performed as described (Reynolds and Fink 2001). We examined the degree of colony ruffling. Biofilms/mats also show differences in colonial expansion (Reynolds and Fink 2001). This parameter was not examined because our controls did not show size differences, and growth differences of alleles complicated interpretation. YPD + 0.3% agar plates were poured and stored at room temperature for 3d. On day three, cells were spread with a toothpick in a circle in the center of each plate. After spreading, plates were wrapped with two sets of parafilm to prevent desiccation and incubated at 32°C for 7d unless otherwise indicated. Each allele was spotted in triplicate. Pictures were taken using the MotiCam ProS5 Plus camera on the eighth day with the Motic Images Plus 3.0 (X64) software set to the following parameters: Exposure, 727.41; Resolution, 1224x1024; Gain, 1; Offset, 0; Enhance, 255 & 0; Gamma, 0.5 & 50; and WhiteBalance Adjust, default. Raw data is available in *File Set S3*.

Images were processed in Python using the Pillow (PIL) (https://pillow.readthedocs.io) and NumPy (https://numpy.org/) libraries. Each image was converted to grayscale using the standard ITU-R 601-2 luma transform (L = 0.299R + 0.587G + 0.114B). Brightness and contrast were standardized by linear rescaling in the 2nd to 98th percentile intensity range to a fixed output range. The same adjustment was applied globally to each image and identically across all images; no local or region-specific adjustments were made. The pipeline used can be found in Code *File Set* S1. Colony ruffling also occurred on standard (YPD + 2% agar) media, see *File Set S1*. Biofilms/mats were scored by visual inspection as smooth, more-wrinkled, or irregular compared to wild type cells and a biofilm-deficient control (*ste12*Δ), which were included for each trial.

To determine the minimum size at which biofilm/mat ruffling was detected, wild-type cells (PC313) were applied by touch to a YPD + 0.3% agar plate with a toothpick. Biofilm/mats were photographed daily for 8d to 11d. Biofilm diameter was measured by ImageJ analysis. Biofilms ≥13 +/- 1 mm diameter (n = 2) showed a ruffled pattern that was scored as wild type (*Fig. S2*). Biofilms/mats formed by spreading cells in a circular pattern (n = 6) also showed same ruffling pattern at the same or smaller diameter as by the touch method.

### Classification of Biofilms/Mats by Pattern Recognition Algorithms

Visual Studio Code and GitHub Copilot were used with the GPT-4.1 Large Language Model (https://openai.com/index/gpt-4-1/). TensorFlow Keras code was used for image classification (https://www.tensorflow.org/tutorials/images/classification). Model training used a deep learning approach based on the MobileNetV2 architecture (https://arxiv.org/abs/1801.04381), a widely used neural network for image classification. The model was trained with >70 images of wild-type biofilms/mats (for ruffled) and >70 images of *ste12* mats (for smooth). All images were used for training except those showing contamination, mis-labelling, or uneven spreading due to mechanical perturbation. For training, images were preprocessed to normalize for height and width differences. To account for variation and prevent bias, the model was trained independently with the same control images. Training was optionally augmented (e.g., by interspersing magnified and rotated images at random) to improve generalization and prevent overfitting the data. While developing the code that produced the final model, training and validation loss / accuracy values were monitored, and training parameters were adjusted until the values followed acceptable trends. Training progress was internally monitored, and the best-preforming model was used for validation.

For validation, the trained model was used to classify each of the 1,018 biofilm/mat images. The algorithm was run in ten separate trials by an automated workflow. Each classification experiment was generated without information from previous runs. The results of each experiment were recorded, and the average values were reported with standard deviation (*Table S8*). Smooth and ruffled predictions were represented as probabilities, which allowed ranking the dataset. The model was compared to visual classification to assess accuracy (*Table S8*). Details of the classification results are available in a sqlite3 database containing the inference results for each image in the dataset (classification_results_20260106_083325.db), and the specific code is on GitHub (https://github.com/CullenSignalingLab/colony-classifier-v1/blob/main/README.md?plain=1).

### β-Galactosidase Assays

β-galactosidase assays were performed as described (Cullen *et al*. 2004) with the following adjustments to assess fMAPK pathway activity in TS mutants. To measure *FRE-lacZ* activity at different temperatures, strains harboring the p*FRE-lacZ* plasmid were patched onto SD-URA semisolid agar media, and plates were incubated at 23°C (or 26°C) for 24h. Cells were incubated in 5mL SD-URA media at 26°C. After 20h of growth, 1.5mL of cells were collected by centrifugation, and resuspended in 3mL SD-URA media on a rotary shaker for 5h at 30°C. Cell densities were measured at A₆₀₀, and cultures were adjusted to an optical density (OD) of ∼1.0. One milliliter of each culture was harvested by centrifugation, and cell pellets were stored at - 80°C. For some experiments, cells were manually collected by a toothpick and resuspended in 1 mL of sterilized water, and A_600_ was measured. Cell cultures were adjusted to A_600_ ∼ 0.6, and 10 μL of OD-adjusted cultures were spotted onto YPD semi-solid agar media for 16h at the temperatures indicated. After 16h, cells were collected in 1 mL of sterilized water, harvested by centrifugation, and stored at -80°C. Cells with the integrated *FUS1-lacZ* reporter were treated similarly except initial patches were made on YPD semisolid agar media.

To perform the ß-galactosidase assays, pellets were unfrozen and resuspended in 100 μL Z-buffer [44.32 mL H_2_O with 5 mL phosphate buffer (0.6 M Na_2_HPO_4_ + 0.4 M Na_2_HPO_4_), 0.5 mL 1 M KCl, 50 μL 1M MgSO_4_, 135 μL β-mercaptoethanol]. Once resuspended, 2 μL of 5% sarkosyl (S) and 2 μL of toluene were added, and the pellets were incubated at 37° C for 30 min with open caps to allow for evaporation of toluene. Then, Z+S buffer containing ortho-Nitrophenyl-β-galactoside (ONPG) was added to the pellets. After color change was observed, reactions were stopped by adding 250 μL of 1M Na_2_CO_3_, and reaction time was recorded. Reaction mixture was centrifuged (13,000 rpm for 3 min) to remove cell extracts, and 200 μL of supernatant was examined at A_420_ to quantitate the yellow color. The A_420_. (1000 X A_420_)/(A_600_ X time) ration was used to calculate Miller units. The average values of two or more biological replicates are reported with error bars showing the standard error of the mean.

### Reverse transcription quantitative PCR Protein

RT-qPCR was performed as described (Chavel *et al*. 2010). Wild-type cells and the *las17-13* mutant were grown in liquid media to mid-log phase. Actin (*ACT1*) and *PGK1* were used as housekeeping control genes.

### Immunoblot Analysis

To detect Sho1 protein levels, strains containing the p*SHO1-GFP* plasmid were grown in SD-URA media for 16 h. Approximately, 500 μL of saturated cell culture was centrifuged, washed, resuspended in 500 μL of sterilized water, and transferred to fresh 10 mL of SD-URA media. Cells were allowed to grow until mid-log phase (A_600_ ∼ 1.0). Five mL of cells were harvested from the mid-log phase cultures for immunoblot analysis. Cells were disrupted, and proteins were enriched by tri-chloroacetic acid precipitation as previously described (Basu *et al*. 2016). Protein samples were subjected to sodium dodecyl sulfate polyacrylamide gel electrophoresis (SDS-PAGE) and transferred to a nitrocellulose membrane (Cat#10600003, Amersham Protran Premium 0.45 μm NC; GE Healthcare Life Sciences). Membranes were blocked in 5% non-fat dried milk for 1 h at room temperature. To visualize Sho1-GFP, membranes were incubated in primary monoclonal mouse anti-GFP antibodies (Cat#11814460001, clones 7.1 and 13.1; Roche) at 1:1,000 dilution. To assess total protein levels, monoclonal mouse anti-Pgk1 antibodies (Life Technologies; Camarillo, CA; Cat #459250) were used at 1:10,000 dilution. Secondary anti-mouse IgG-HRP (Cat#1706516; Bio-Rad, Inc.) and goat anti-rabbit IgG-HRP (Cat#115-035-003; Jackson ImmnunoResearch Laboratories) were used to detect the primary antibodies. Primary antibody incubations were performed at 4° C for 16 h, and secondary antibody incubations were performed at 22° C for 1 h. Immunoblots were visualized by Gel Doc XR Imaging System (Bio-Rad, Inc.), after addition of Chemiluminescent HRP substrate for chemiluminescent Westerns (Radiance Plus Substrate, Azure Biosystems).

For phospho-MAP kinase levels, protein sample preparation was performed as above. Membranes were probed with rabbit polyclonal p44/42 antibodies (Cell Signaling Technology, Danvers, MA; Cat #4370) diluted 1:10,000 in 5% bovine serum albumin (BSA) to detect P∼MAP kinases, P∼Fus3 and P∼Kss1. Monoclonal mouse anti-Pgk1 antibodies (Life Technologies; Camarillo, CA; Cat #459250) were used at 1:10,000 dilution as a control for total protein levels. Secondary anti-mouse IgG-HRP (Cat#1706516; Bio-Rad, Inc.) and goat anti-rabbit IgG-HRP (Cat#115-035-003; Jackson ImmnunoResearch Laboratories) were used to detect the primary antibodies. The nitrocellulose membrane was blocked with 5% non-fat dried milk for Pgk1 antibody or 5% BSA for the p44/42 antibody for 1 h prior to antibody incubation. Densitometric analysis was performed with Image Lab Software (Bio-Rad https://www.bio-rad.com/en-us/product/image-lab-software?ID=KRE6P5E8Z). Exposures in the linear range were used for measuring band intensities. Background subtraction was performed according to the guidelines provided by the manufacturer. Band intensities of phospho-proteins were normalized against total protein levels based on Pgk1 band intensity. Wild-type levels were normalized to Pgk1 levels and set to a value of one. Other normalized values were adjusted accordingly.

### Halo assays to assess mating pathway activity

Halo assays were performed as described (Julius *et al*. 1983) at the same temperatures (30°C and 32°C) as fMAPK assays. To determine the percentage of polarized cells in response to mating factor (shmoos), wild-type cells and a panel of alleles were grown to mid-log phase. Cells were examined by microscopy at 100X.

### Microscopy

Scanning electron microscopy was performed as described (Basu *et al*. 2016) with modifications described in (Prabhakar *et al*. 2020). Differential interference contrast (DIC) microscopy was performed as described (Basu *et al*. 2016) with a Zeiss Axioplan 2 microscope. Time-lapse microscopy was performed as described (Prabhakar *et al*. 2020).

### Protein Localization by Fluorescence Microscopy

To visualize Sho1-GFP, strains containing p*SHO1-GFP* plasmid were grown in SD-URA for 16 h at 30°C, sub-cultured into SD-URA media (∼500 μL saturated culture in 10 mL fresh media) and grown until mid-log phase (A_600_ ∼ 1.0) before harvesting cells for fluorescence microscopy. 1 mL of cells were centrifuged, washed twice with sterilized water, and resuspended in 500 μL of sterilized water. Differential interference contrast (DIC) filter was used to visualize cells, and fluorescein isothiocyanate (FITC) filter was used to visualize GFP. An Axioplan 2 fluorescence microscope (Zeiss) was used with a Plan-Apochromat 100×/1.4 (oil) objective (N.A. 1.4; cover slip 0.17) with the Axiocam MRm camera (Zeiss). Images were analyzed using Axiovision 4.4 software (Zeiss). For some experiments, strains were patched onto SD-URA semi-solid agar media. Plates were incubated at 30°C for 16 h, and cells were examined directly on plates with coverslips at 100X magnification. The localization of proteins driven by the strong *GAL1-10* promoter were determined in cells grown in S-Gal-URA media. To measure circularity, DIC images were imported in ImageJ (https://imagej.nih.gov) and the length and width of cells were measured by drawing a line across the cells at their long and short axes and using the measure tool to determine length. Length and width measurements were exported in Excel, and length/width was used to determine circularity.

### Bioinformatics Analysis

Some models were made by Adobe Illustrator. DNA comparisons were performed by Benchling (https://www.benchling.com/). Graphs were made with Excel. Alpha fold prediction programs (Jumper *et al*. 2021) were performed using ChimeraX (https://www.cgl.ucsf.edu/chimerax/) as described (Goddard *et al*. 2018; Pettersen *et al*. 2021; Meng *et al*. 2023). Statistical analysis was performed in Minitab for ANOVA analyses. Some statistical analyses were performed with Prism (https://www.graphpad.com/features). Brightness/contrast adjustments were made to some photos with ImageJ (https://github.com/fiji/fiji). Images generated from the BioRad Gel Documentation system as (.scn) were converted in batch to (.tiff) files using a macro script generated by CHATGPT (https://chatgpt.com/) by ImageJ analysis (FIJI), the script can be found in Code File S2.

### Data availability statement

Strains, plasmids, and the Sigma ts collection are available upon request to.

### Use of AI Software

AI software was used for data analysis (see Materials & Methods). CHAT GPT (https://chatgpt.com/) and Anthropic (https://claude.ai/new) were used for editorial purposes to help with sentence structure, grammar, and logic flow.

## ABBREVIATIONS

3-ATA: 3-aminotriazole
5-FOA: 5-floroorotic acid
DIC: Differential Interference Contrast
DMSO: dimethylsulfoxide
G-protein: GTP binding protein
GFP: green fluorescent protein
GPCR: GTP binding-protein coupled receptor
IB: immunoblot
IP: immunoprecipitation
kDa: kilodalton
MAPK: mitogen activated protein kinase
ND: not determined
OD: optical density
ONPG: ortho-Nitrophenyl-β-galactosidase
ORF: open reading frame
PAK: p21-activated kinase
PCR: polymerase chain reaction
PH: pleckstrin homology
PKA: protein kinase A
Rho: Ras homology
ROI: region of interest
RTG: retrograde-to-nucleus
SDS-PAGE: sodium dodecyl sulfate polyacrylamide gel electrophoresis
ts: temperature sensitive
WASP: Wiskott-Aldrich Syndrome Protein
WASH: Wiskott-Aldrich Syndrome Protein Homolog
WT: wild type.

The work was supported by an award from the NIH (to PJC GM098629).

## ACKNOWLEDGEMENTS

Thanks to Daniel Lew (Duke University), Alan Davidson (University of Toronto), Peter Walter (University of California San Francisco), David Eide (University of Wisconsin at Madison), Zhengchang Liu (University of New Orleans), Michael Breitenbach (University of Salzburg, Salzburg Austria), Hiten Madhani (University of California San Francisco), and P. Pryciak (U MASS, Amherst, MA) for providing reagents. Thanks to Leah Cowen (University of Toronto) and Jeremy Thorner (University of California Berkeley) for helpful suggestions. Thanks to Somdatta Maiti, Shohely Anonna, Ariana Johnson, Adriana Victoria Acosta Ortiz, Chengchen Gao, and Antonio DeVincentis for help with the screen. Thanks to Sarah Walker (University at Buffalo) for helpful comments and sharing equipment. Thanks to Katherine (Kay) Yevzerov for help with data analysis. Thanks to Animesh Bote for helping with biofilm/mat raw image processing. Thanks to laboratory members for reading the manuscript and providing suggestions.

## Author contributions

ANP designed and performed experiments, analyzed data, wrote, and edited the paper; AP, performed experiments and analyzed data; ZL, performed experiments and provided reagents and technical assistance; ASD performed experiments, analyzed the data, and edited the paper; DW performed experiments; DC, SB, and HF provided reagents and technical assistance; ML, performed experiments; JO, performed experiments; RMR, developed bioinformatics systems and analyzed data; BA and CB provided reagents, suggestions, technical assistance and expertise; PJC, designed and performed experiments, wrote and edited the paper, and obtained funding for the study.

## SUPPLEMENTAL MATERIALS

### SUPPLEMENTAL FIGURE LEGENDS

**Figure S1.**
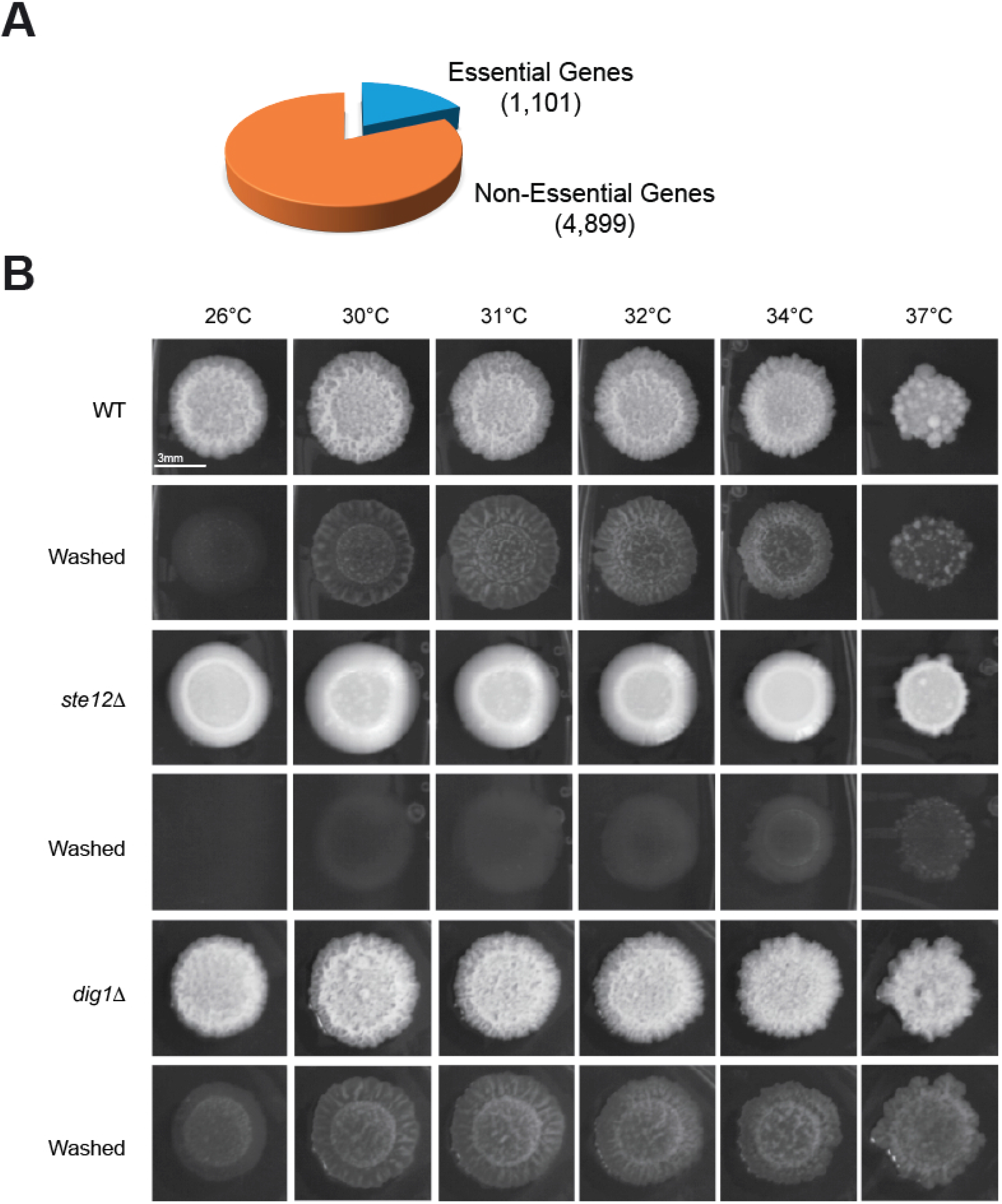
Essential genes in yeast and the temperature dependence of filamentous growth. **A)** Pie chart of the relative essential to nonessential genes in yeast. **B)** The indicated strains were incubated at the indicated temperatures for 5d, photographed, washed, and photographed again. Bar, 3 mm.

**Figure S2.**
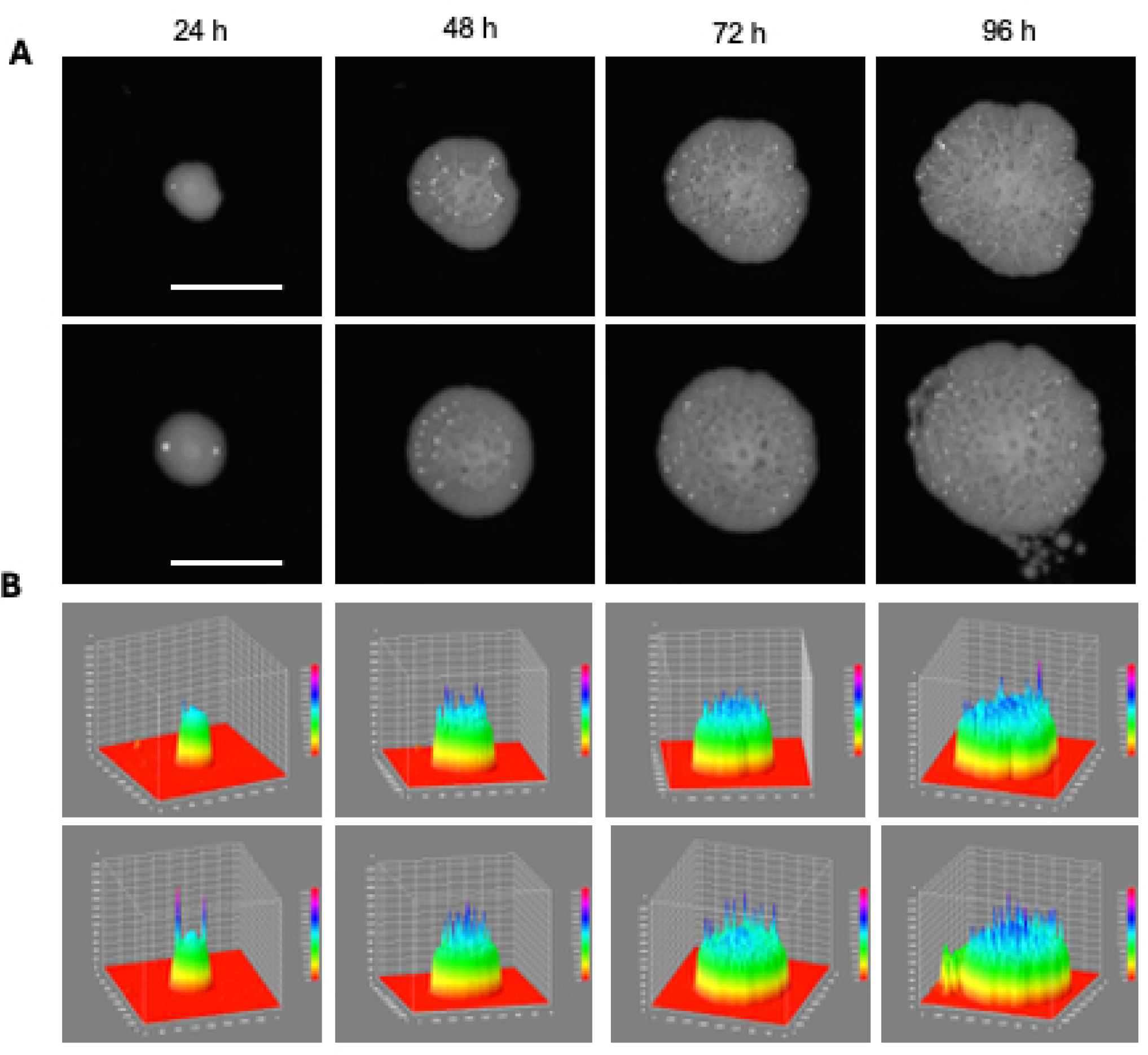
Determination of the minimum size for detecting biofilm/mat ruffling. **A)** Wild-type cells (PC313) were inoculated at the center of YPD + 0.3% agar plates with a sterile toothpick. Colony ruffling was monitored over a 96-h period to assess the ruffled patterns at 32°C. Two biological replicates are shown. Bar, 12 mm. **B)** Ruffled patterns in panel *S2A* by ImageJ viewed as a surface plot using the plugin Interactive 3D Surface Plot (https://imagej.net/ij/plugins/surface-plot-3d.html).

**Figure S3.**
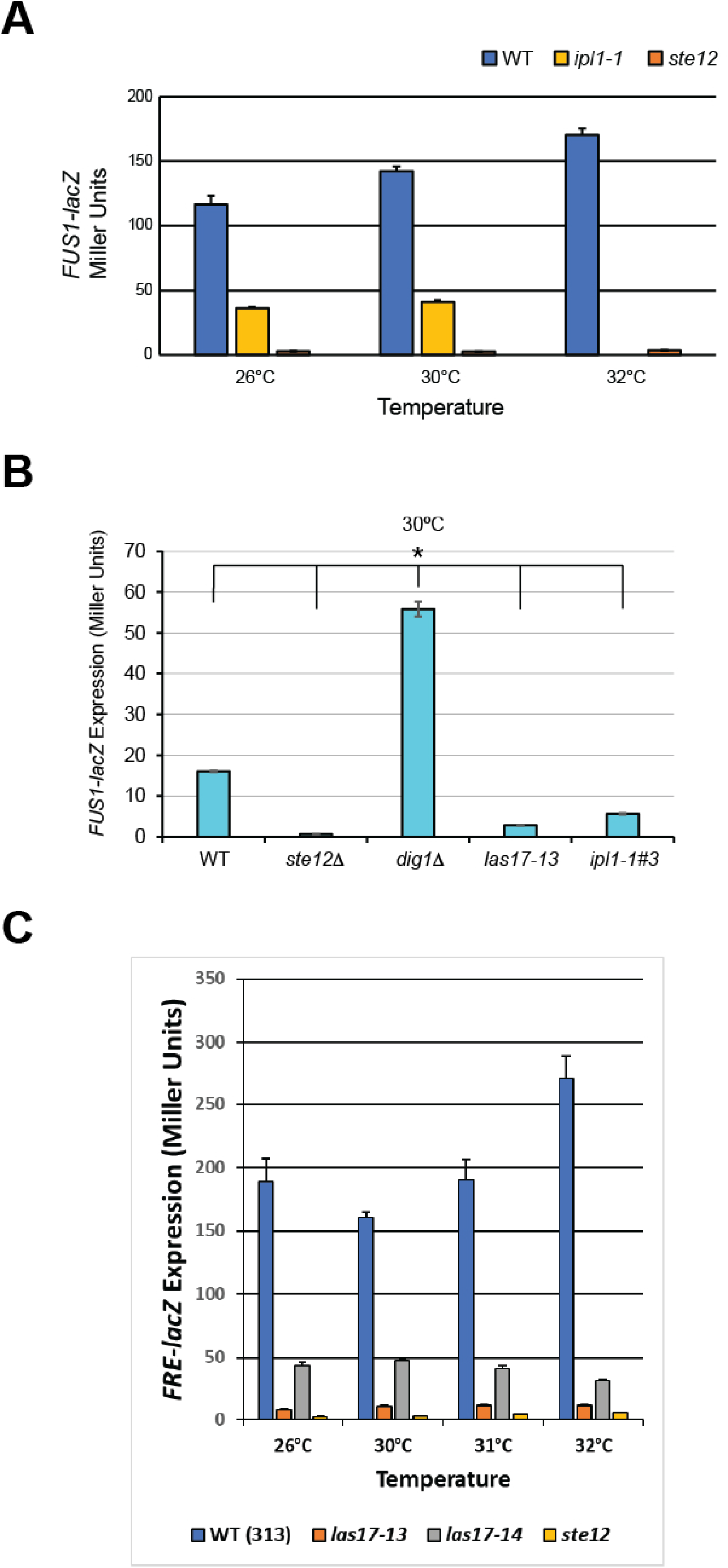
Evaluating ts alleles for variation in fMAPK pathway activity. **A)** β-galactosidase assays. *FRE-lacZ* reporter activity in the indicated strains at 26°C, 30°C, and 32°C. Error bars represent the standard error of the mean. **B)** β-galactosidase assays. *FUS1-lacZ* reporter activity in the indicated strains at 30°C. Experiments were performed in three biological replicates. Data were analyzed by one-way ANOVA followed by a Tukey’s pairwise comparison test to generate p-values. Error bars denote standard error of the mean. Asterisk denotes difference compared to WT and p-value < 0.05. **C)** β-galactosidase assays. fMAPK pathway activity was assessed by the *FRE-lacZ* reporter. β-Galactosidase assays were performed on the indicated strains as described. Miller units are reported relative to WT values. Experiments were performed in three biological replicates. Data were analyzed by one-way ANOVA followed by a Tukey’s pairwise comparison test to generate p-values. Error bars denote standard error of the mean.

**Figure S4.**
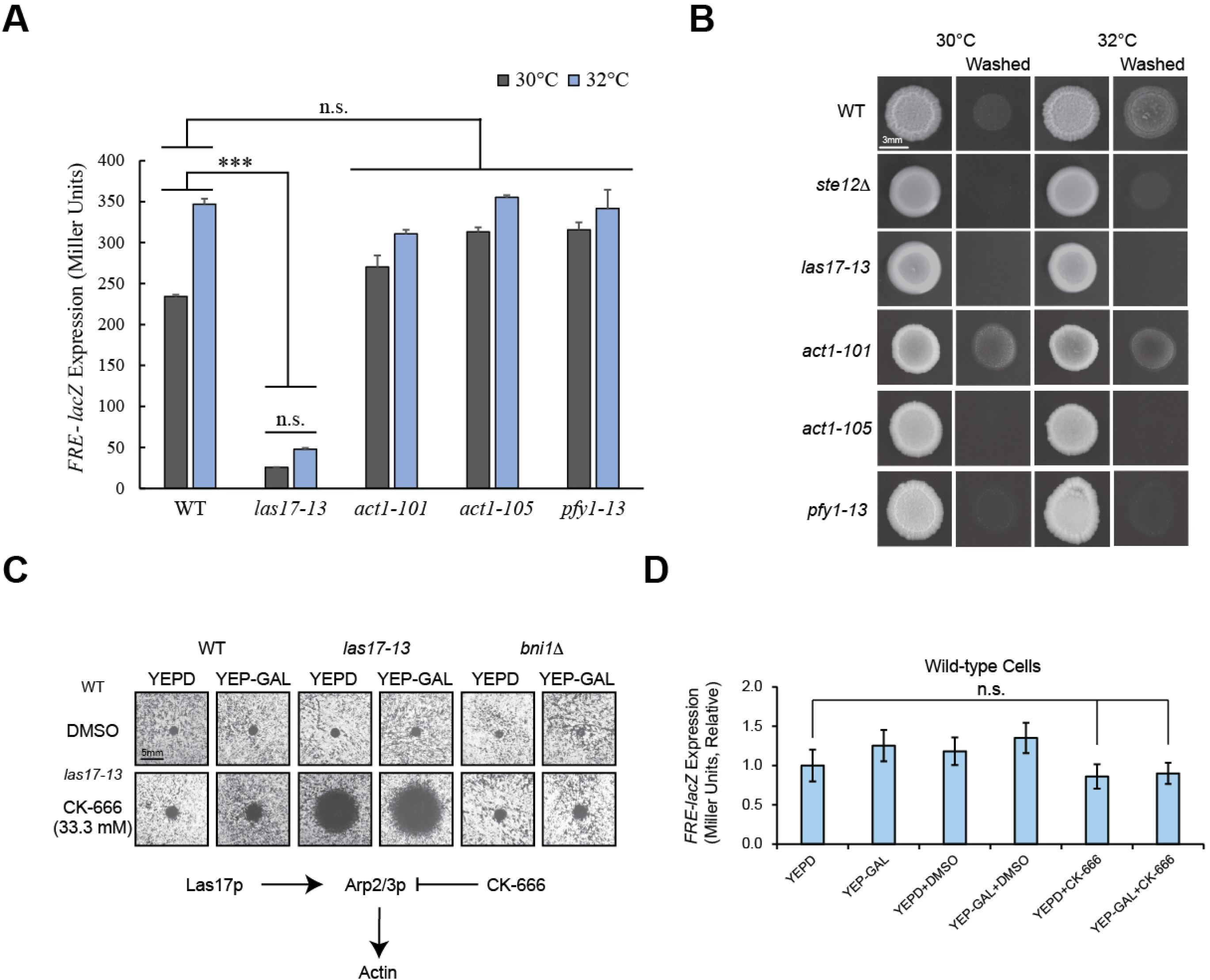
Role of mutants defective for actin function in filamentous growth and fMAPK pathway activity. **A)** β-galactosidase assays. Strains harboring the p*FRE-lacZ* reporter were spotted on YPD media, and plates were incubated at indicated temperatures for 16 h. Cells were scraped from plates using toothpicks into sterile water, and ß-galactosidase assays were performed in three biological replicates. Data were analyzed by one-way ANOVA followed by a Tukey’s pairwise comparison test to generate p-values. Asterisk denotes difference compared to WT and p-value<0.05. Error bars denote standard error of the mean. **B)** The plate-washing assay. WT cells (PC313), *ste12*Δ mutant, and conditional alleles *las17-13*, *act1-105, act1-101,* and *pfy1-13* were spotted on YPD media. Plates were incubated at the indicated temperatures for 3 d. To reveal invasive scars, colonies were washed under a stream of water, and pictures were taken before and after washing. Bar, 3 mm. **C)** Growth inhibition by CK-666. WT cells (PC313), and the *bni1*Δ and *las17-13* mutants were top-spread on YPD and YEP-GAL media. 10 uL of dimethylsulfoxide (DMSO) or 33.3 mM CK-666 was spotted in the center of the plate. Plates were incubated for 2d at 30°C. Bar, 5 mm. **D)** ß-galactosidase assays in wild-type cells treated with CK-666. Wild-type (PC313) cells harboring the p*FRE-LacZ* plasmid were grown in YPD and YP-GAL media at 32°C and treated with DMSO and 33.3 mM CK-666. Experiments were performed in three independent replicates. Wild-type values were set to a value of one and other values were adjusted accordingly. Error bars denote standard error of the mean. Data were analyzed by one-way ANOVA followed by a Tukey’s pairwise comparison test to generate p-values. None of the values showed a statistically significant difference compared to wild type (ns, not significant).

**Figure S5.**
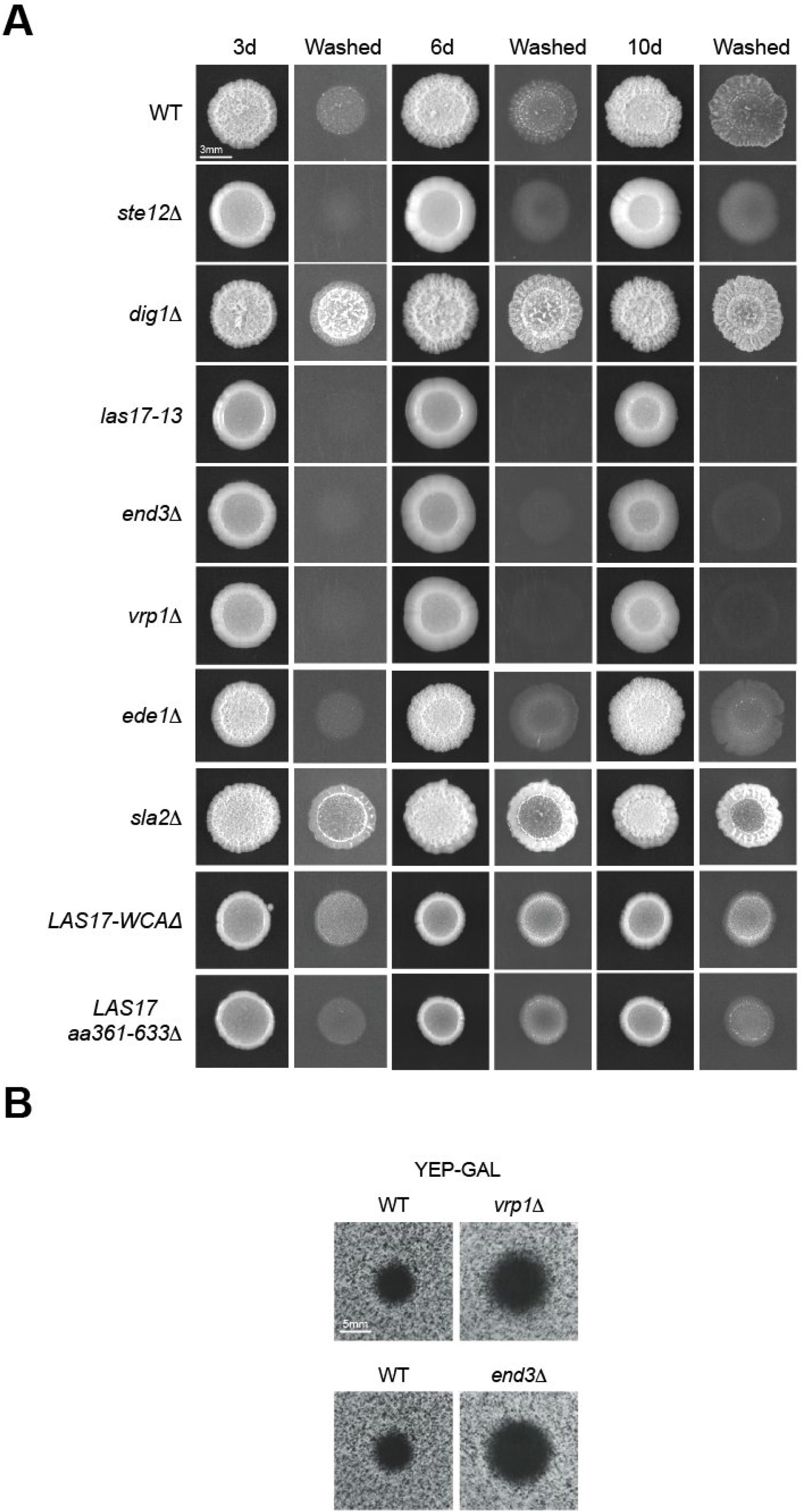
Role of proteins that control endocytosis in regulating filamentous growth. **A)** The WT (PC313), *ste12*Δ (PC539), and *dig1*Δ (PC3039) control strains were spotted on YPD semi-solid agar media along with endocytosis mutants *end3*Δ (PC7920), *vrp1*Δ (PC7863), *ede1*Δ (PC7887), *sla2*Δ (PC7922), and two separate isolates of the *LAS17-WCA*Δ mutant. Plates were incubated at 30°C for 3d, 6d, and 10d. On the indicated days, colonies were photographed, washed under a stream of water, and photographed again. Bar, 3 mm. **B)** Sensitivity of the indicated mutants to CK-666 in YPD and YP-GAL media at 30°C was examined. Bar, 5 mm.

**Figure S6.**
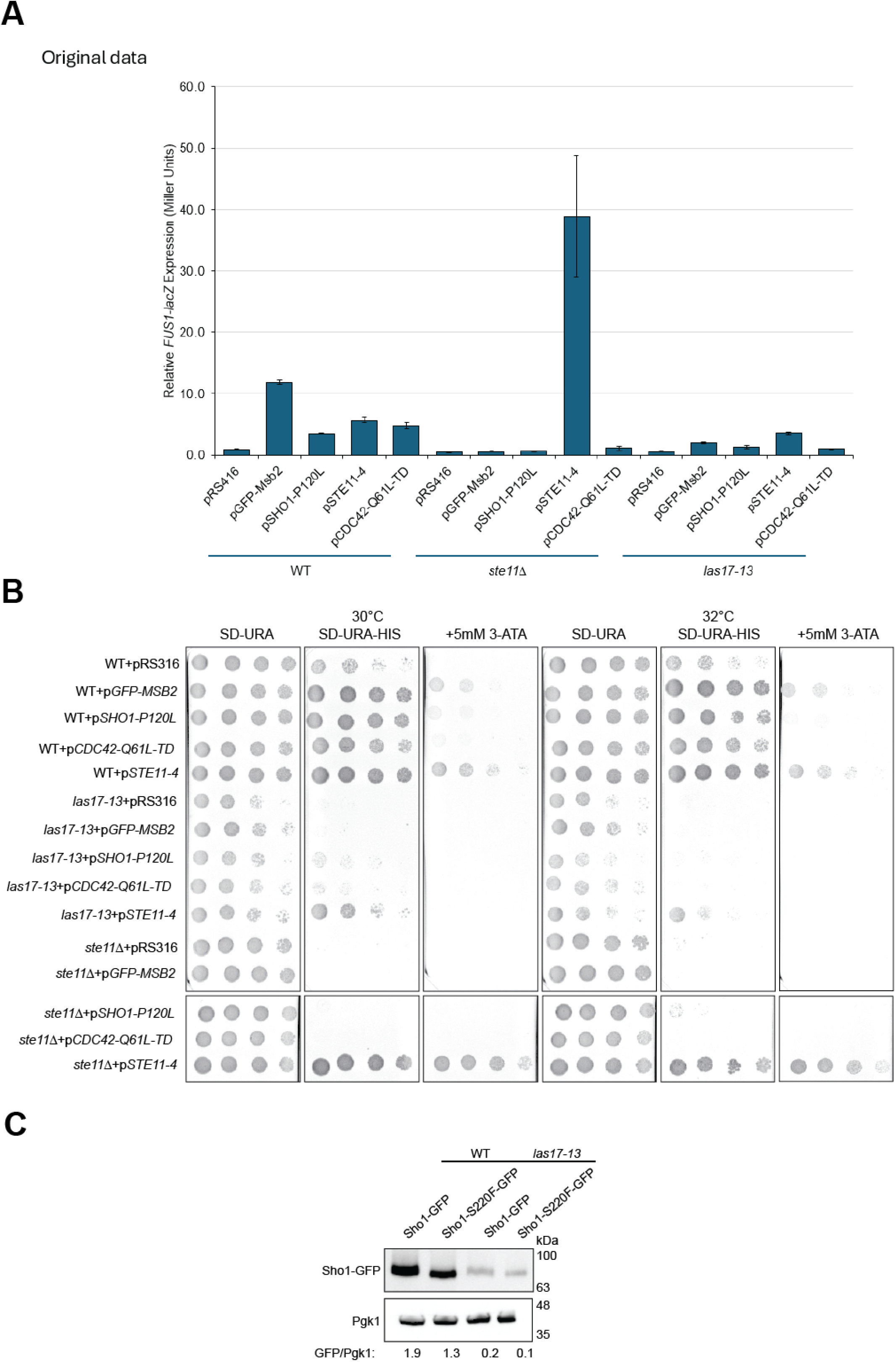
Genetic suppression analysis of the *las17-13* and other mutants in combination with alleles that hyperactivate the fMAPK pathway. A) Bar graph of *FUS1-lacZ* activity in the indicated strains lacking *STE4.* Experiments were performed in duplicate, and error bars show the standard deviation between samples. B) Activity of the *FUS1-HIS3* growth reporter. Controls at left. ATA, aminotriazole resistance. C) IB of pSHO1^S220F^-GFP. levels in wild-type cells and the *las17-13* mutant. See Fig. 4E for details. Antibodies to Pgk1 were used as a loading control. Relative band intensities are shown.

**Figure S7.**
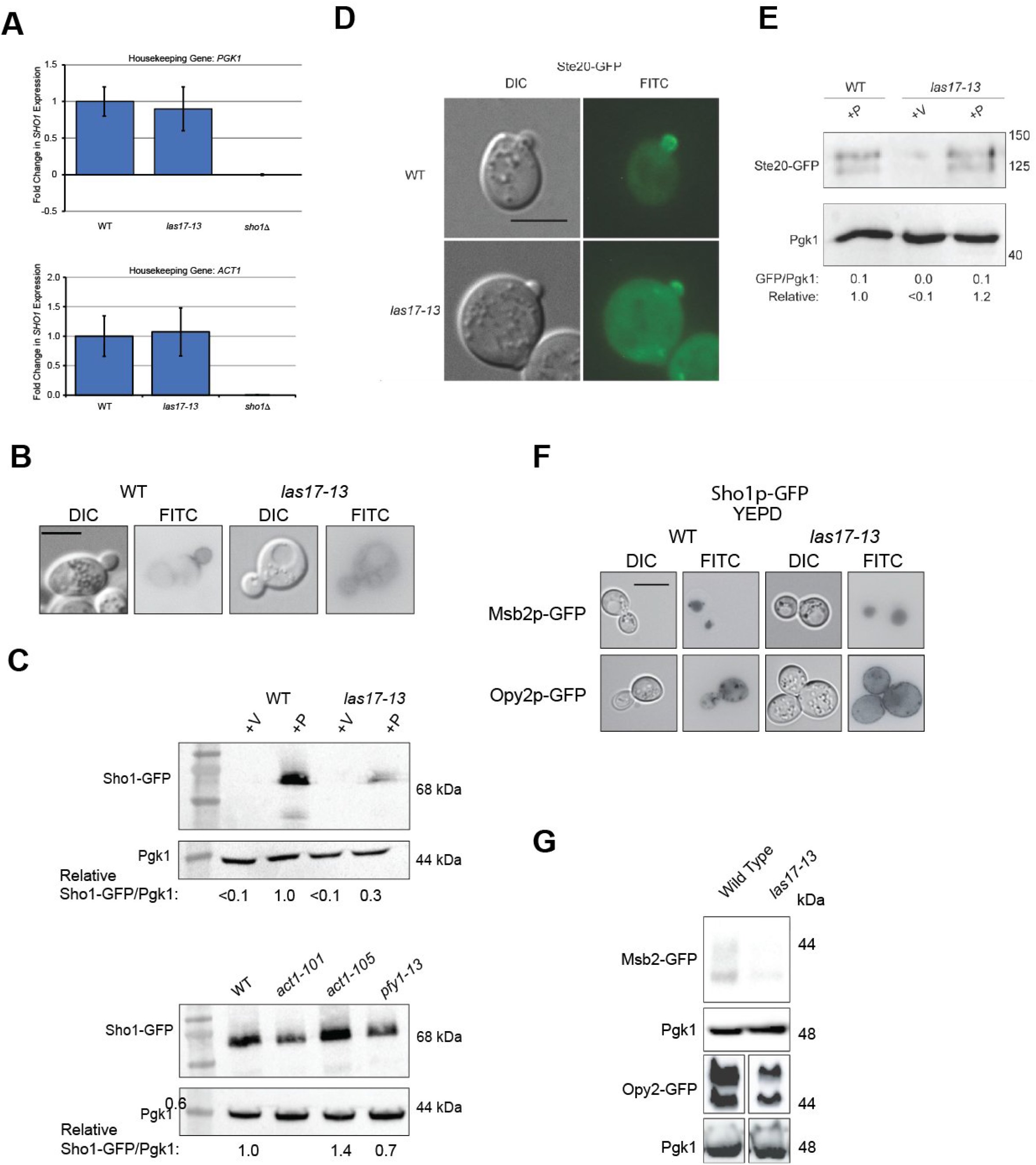
Exploring regulatory features of fMAPK pathway components. **A)** RNA levels of *SHO1* relative to *PGK1* (top panel) or *ACT1* (bottom panels) in wild-type cells and the *las17-13* mutant. **B)** Top panels, the localization of pSho1-GFP in wild-type cells and the *las17-13* mutant at 32°C. Bar, 5 microns. **C)** Immunoblot analysis for Sho1-GFP levels in actin cytoskeleton mutants, *act1-101*, *act1-105*, and *pfy1-13* introduced by plasmid transformation. **D)** Wild-type cells [WT (PC313)] and the *las17-13* mutant containing *STE20-GFP* on a plasmid (PC4394) were observed at 100X magnification using the DIC channel to visualize cells and the FITC channels to visualize localization of Ste20 tagged with GFP protein. Scale bar, 5 microns. **E)** Immunoblot analysis using antibodies to visualize Ste20-GFP, and Pgk1 as control for total protein levels. +V denotes strain with pRS316 (PC2207) as a control, and +P denotes strains containing p*STE20-GFP.* **F)** Localization of pMsb2-GFP and pOpy2-GFP in wild-type cells (WT, PC313) and the *las17-13* mutant grown in YEPD. For each strain, DIC and FITC channels are shown. Bar, 5 microns. **G)** Levels of Msb2-GFP and Opy2-GFP in wild-type cells and the *las17-13* mutant assessed by immunoblot analysis, with blots measuring Pgk1 levels as a loading control. Molecular weights (kDa) are indicated.

**Figure S8.**
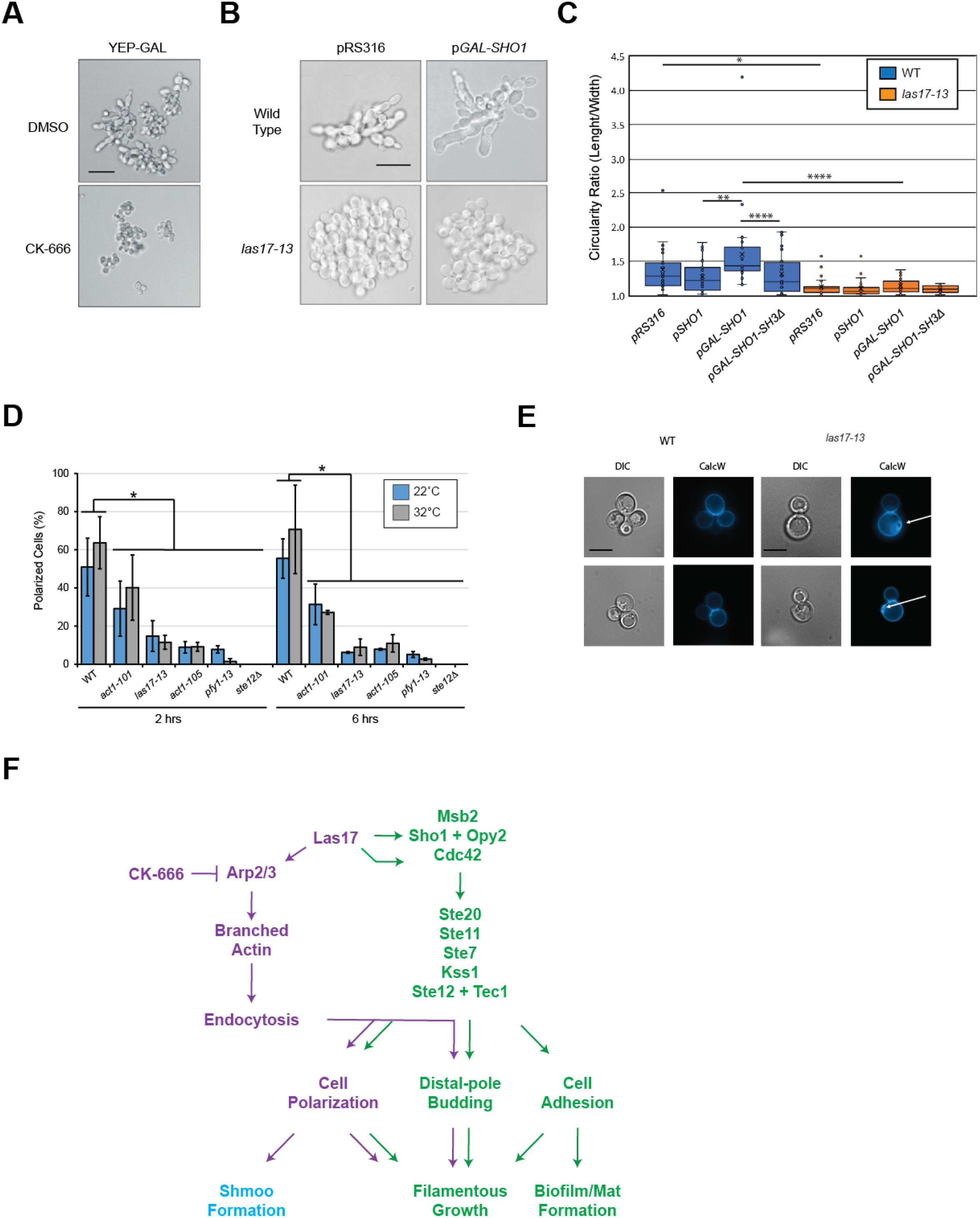
Global roles for Las17 in filamentous growth regulation and cell polarization in yeast. **A)** Wild-type (PC 313) cells were inoculated in 5 mL of YPD media and grown at 30°C till they reached saturation (∼16 h). Cells were diluted to 0.03 OD600 in fresh YPD media, and 200 μL of diluted culture was added to 96-well plate. 5 μL of CK-666 (Stock Conc. 3.3 mM) was added to the 200 μL culture. Plates were incubated at 30°C for 48 hrs. Cells were looked at after 48 h at 100X magnification using the DIC filter to observe the formation of filaments. To observe the effect of CK-666 on YEP-GAL-induced filamentous growth, 5 mL of saturated culture of WT cells was pelleted, washed twice with 1 mL of sterilized water, and transferred to fresh 5 mL of YEP-GAL media with 12 μL of 33 mM CK-666. Cells were grown for 6 h and observed under the microscope at 100X using DIC filter. Bar, 5 microns. **B)** Las17 is required for Sho1-dependent cell polarization. Wild-type cells (PC313) and the *las17-13*, *act1-105*, and *pfy1-13* mutants containing p*RS316* or p*GAL-SHO1-GFP* plasmids were grown in S-GAL-URA media at 32°C visualized using the DIC filter at 100X magnification to observe *SHO1*-dependent hyperpolarized morphology. Bar, 5 microns. **C)** Cell circularity (Length/Width) was measured in wild-type cells (PC313) and the *las17-13* mutant incubated at 32°C harboring p*RS316*, p*SHO1-GFP*, p*GAL-SHO1-GFP,* and p*GAL-SHO1-SH3*Δ*-GFP* plasmids. Box plot is shown, n =25 cells. Data were analyzed by one-way ANOVA followed by a Tukey’s pairwise comparison test to generate p-values. Asterisk denotes the difference compared to WT; p-value < 0.05. **D)** Las17 regulates polarized growth during mating. Wild-type (PC313) and *las17-13* cells harboring p*SHO1-GFP* were inoculated in 5 mL of SD-URA media and grown at 30°C till they reached saturation (∼16 h). Cells were diluted to 0.03 OD600 in fresh YPD media and grown till they reached A_600_ ∼ 1.0. 1 mL of culture was removed and treated with 10 μL of α-factor (stock conc. 1 mg/mL). Cells were quantitated for shmoo formation. **E)** Calcofluor White staining of wild-type cells and the *las17-13* mutant grown at 30°C. Bar, 5 microns. **F)** Model for the regulatory inputs of Las17 in the fMAPK pathway.

**Figure S9.**
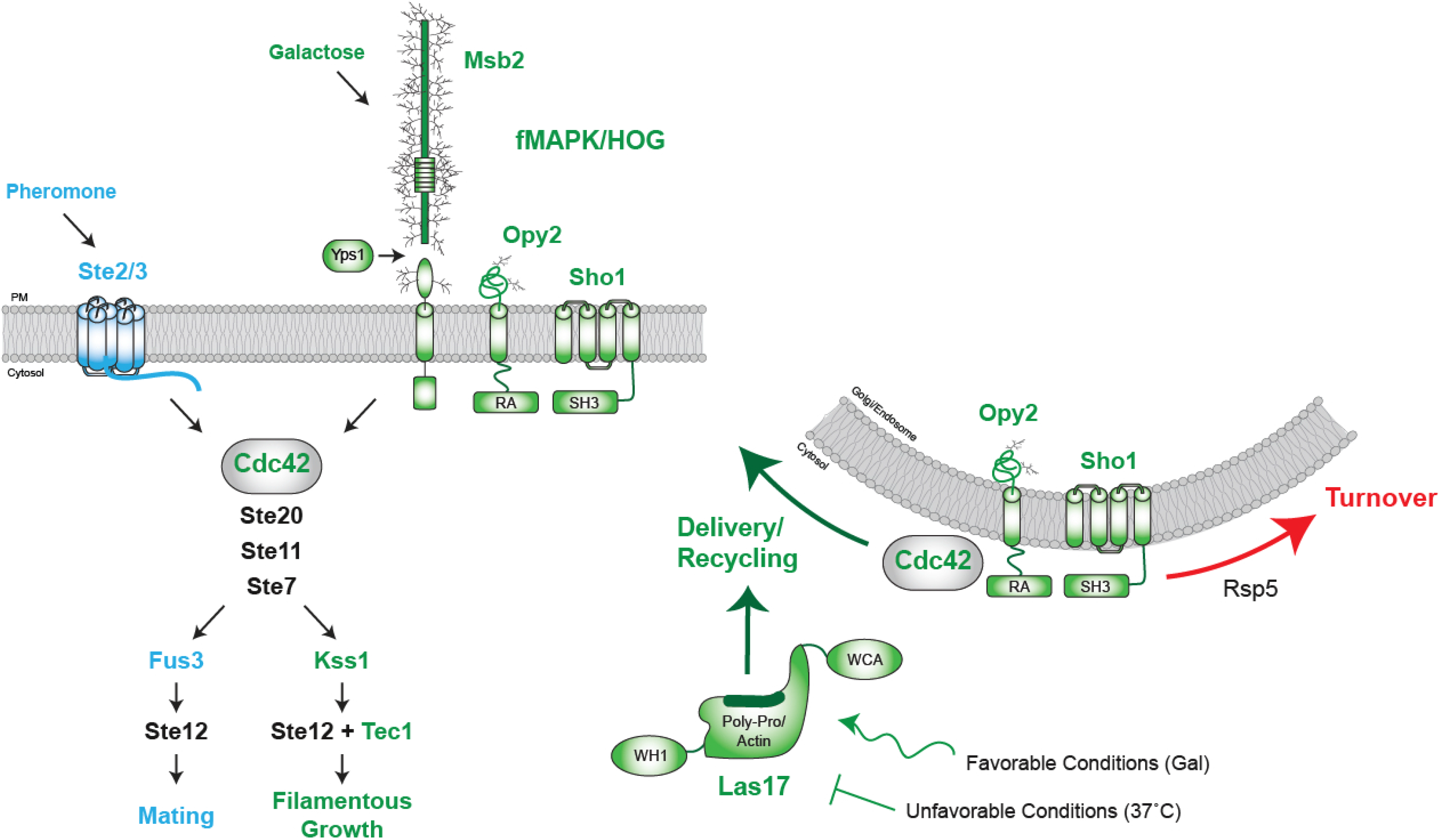
Model for how Las17 regulates the fMAPK pathway. Two MAPK pathways share common components. Mating pheromone is detected by a G-protein coupled receptor (GPCR, Ste2 and Ste3), which functions through a specific MAP kinase (Fus3) to allow mating (blue). Glucose limitation induces a tripartite complex of the mucin Msb2, tetra-span protein, Sho1, and the cysteine-rich protein, Opy2, which function through a different MAP kinase (Kss1) and transcription factor (Tec1) to induce filamentous growth (green). The two pathways share common components (Cdc42, Ste20, Ste11, Ste7, and Ste12), although Cdc42 levels are singularly important for fMAPK pathway activation (Gonzalez and Cullen 2022). Las17 promotes the cell-surface delivery and/or recycling of Msb2, Sho1, Opy2, and Cdc42, which presumably occurs from internal compartments. This action delays turnover of the proteins in the trafficking pathway. The levels of Las17 may be regulated to sustain fMAPK pathway activity. Some determinants of Sho1 trafficking have been mapped to regions in the C-terminal cytosolic region including the SH3 domain.

### SUPPLEMENTAL RAW DATA FILE SETS

*File Set S1.* The raw data for the plate-washing assay can be found at the following link: https://buffalo.app.box.com/folder/169493491171. Mutant alleles and controls were spotted in duplicate onto YPD media and incubated at 30°C, 32°C, and 34°C for 4d. Plates were photographed, washed in a stream of water, and photographed again. The key for each plate can be found in the associated excel spreadsheet.

*File Set S2.* The raw data containing the single cell assay can be found at the following link: https://buffalo.app.box.com/folder/161505250854. The indicated strains were spotted onto S-GLU media. Plates were incubated for 16h and representative examples of cells were photographed. Cells were incubated at 32°C and for some alleles also at 30°C. The folder is searchable by keyword.

*File Set S3.* The raw data containing the biofilm/mat data can be found at the following link: https://buffalo.app.box.com/folder/174116431845. The indicated strains were patched in at least three biological replicates onto YPD + 0.3% agar media for 7d at 32°C. Plates were photographed.

The folder can be searched by keyword.

