## Supplementary material for "Exploring roles for essential proteins in yeast filamentous growth identifies the WASP homolog Las17 as a regulator of the Cdc42-dependent fMAPK pathway": Table S1

**Table S1. Yeast strains used in the study.**

| Strains | Genotype^a^ | Reference |
| --- | --- | --- |
| PC312 | *MAT*α *ura3-52* | (Gimeno *et al.* 1992) |
| PC313 | *MAT***a** *ura3-52* | (Gimeno *et al.* 1992) |
| PC344 | *MAT*α *ura3-52 / MAT***a** *ura3-52* | (Cullen *et al.* 2004) |
| PC538 | *MAT***a** *ste4 FUS1-lacZ FUS1-HIS3 ura3-52* | (Cullen *et al.* 2004) |
| PC539 | *MAT***a** *ste4 FUS1-lacZ FUS1-HIS3 ura3-52 ste12::KlURA3* | (Cullen and Sprague 2000) |
| PC1160 | *MAT***a** *ste4 FUS1-lacZ FUS1-HIS3 ura3-52 bni1::URA3* | (Cullen and Sprague 2000) |
| PC611 | *MAT***a** *ste4 FUS1-lacZ FUS1-HIS3 ura3-52 ste11::URA3* | (Cullen *et al.* 2004) |
| PC6222 | *MAT***a** *ste4 FUS1-lacZ FUS1-HIS3 ura3-52 ras2::HYG* | (Chavel *et al.* 2014) |
| BY4741^c^ | *MAT***a** *ura3*Δ*0 leu2*Δ*0 met15*Δ*0 his3*Δ*1* | (Winzeler *et al.* 1999) |
| PC6032 | *MAT***a** *ste4 FUS1-lacZ FUS1-HIS3 ura3-52 ire1::URA3* | (Adhikari *et al.* 2015b) |
| PC3695 | *MAT***a** *ste4 FUS1-lacZ FUS1-HIS3 ura3-52 rtg1::URA3* | (Chavel *et al.* 2014) |
| PC2710 | *MAT***a** *ste4 FUS1-lacZ FUS1-HIS3 ura3-52 NAT::cdc12-6* | (Chavel *et al.* 2014) |
| PC3039 | *MAT***a** *ste4 FUS1-lacZ FUS1-HIS3 ura3-52 MSB2-HA dig1::KlURA3* | (Chavel *et al.* 2014) |
| PC7365 | *MAT***a** *ura3-52 CDC3-mCHERRY::HYG* | (Prabhakar *et al.* 2020) |
| SigmaTS^b^ | *MAT***a** *ura3-52 las17-1::KanMX6* | This study |
| SigmaTS^b^ | *MAT***a** *ura3-52 las17-13::KanMX6* | This study |
| SigmaTS^b^ | *MAT***a** *ura3-52 las17-14::KanMX6* | This study |
| SigmaTS^b^ | *MAT***a** *ura3-52 act1-101::KanMX6* | This study |
| SigmaTS^b^ | *MAT***a** *ura3-52 act1-105::KanMX6* | This study |
| SigmaTS^b^ | *MAT***a** *ura3-52 pfy1-13::KanMX6* | This study |
| PC7920 | *MAT***a** *ura3-52 end3::KanMX6* | This study |
| PC7693 | *MAT***a** *ura3-52 leu2* | This study |
| PC7863 | *MAT***a** *ura3-52 vrp1::KanMX6* | This study |
| PC7921 | *MAT***a** *ura3-52 myo3::KanMX6* | This study |
| PC7861 | *MAT***a** *ura3-52 myo5::HYG* | This study |
| PC7922 | *MAT***a** *ura3-52 sla2::KanMX6* | This study |
| PC7858 | *MAT***a** *ura3-52 ede1::KanMX6* | This study |
| PC7853 | *MAT***a** *ura3-52 LAS17-WCA*Δ*::KanMX6* | This study |
| PC7711 | *MAT***a** *ura3-52 leu2 his3* | This study |
| ATP101 | *MAT***a** *ura3-52 las17aa361-633*Δ*::KanMX6* | This study |

a. All strains are in the ∑1278b background unless otherwise indicated.
b. For the strains made in the SigmaTS collection refer to *Table S4*.
c. Strain background *S288c*.
