## Supplementary material for "Exploring roles for essential proteins in yeast filamentous growth identifies the WASP homolog Las17 as a regulator of the Cdc42-dependent fMAPK pathway": Table S2

**Table S2. Plasmids used in the study.**

| Location | Name | Description/Markers | Reference |
| --- | --- | --- | --- |
| PC1308 | *pFRE-lacZ* | 2 micron/URA3 | (Madhani and Fink 1997) |
| PC2560 | pGFP-2XPH-PLCδ | CEN/URA | (Stefan *et al.* 2002) |
| PC1964 | pSHO1-GFP | CEN/URA3 | (Marles *et al.* 2004) |
| PC1715 | pSHO1^P120L^-GFP | CEN/URA3 | (Vadaie *et al.* 2008) |
| PC1717 | pSHO1^S220F^-GFP | CEN/URA3 | (Vadaie *et al.* 2008) |
| PC7143 | pCDC12-GFP::KanMX6 | LEU2/CAN KanMX6 | This study |
| PC7450 | pGPD1-LAS17-FLAG | URA3/CEN | (Weber *et al.* 2021) |
| PC1882 | pGAL-SHO1-GFP | URA3/CEN/KanMX6 | (Pitoniak *et al.* 2015) |
| PC2207 | pRS316 | URA3/CEN | (Sikorski and Hieter 1989) |
| PC6454 | pGFP-Cdc42 | URA3/CEN | (Gonzalez and Cullen 2022) |
| PC3944 | pOpy2-GFP | URA3/CEN | (Karunanithi and Cullen 2012) |
| PC2582 | pHA-Msb2-GFP | URA3/CEN/KanMX6 | (Vadaie *et al.* 2008) |
| PC7361 | pSho1-GFP::KanMX6 | URA3/CEN | $(Prabhakar et al. 2020) |
| PC1616 | pSho1-GFP-SH3Δ | URA3/CEN | (Marles *et al.* 2004) |
| PC6454 | pGFP-Cdc42 | CEN/URA | (Gonzalez and Cullen 2022) |
| PC7458 | pGFP-Cdc42^Q61L^ | CEN/URA | (Gonzalez and Cullen 2022) |
| PC7654 | pGFP-Cdc42^Q61L+TD^ | CEN/URA | (Gonzalez and Cullen 2022) |
| PC1881 | pGAL-SHO1-SH3Δ-GFP | CEN/URA/KanMX6 | This study |
